# Geoclimatic oscillations and ancient reciprocal adaptive introgression shape the evolutionary trajectories of threatened *Coilia*

**DOI:** 10.64898/2026.02.19.706424

**Authors:** Zhenqiang Fu, Hao Wang, Yidong Feng, Yan Hu, Zhongwei Sun, Zhanyuan Gao, Weiwei Zhang, Xuanguang Liang, Qinglong Chen, Shan Liu, Baolong Bao, Jianguo Lu

**Author notes:** Corresponding author: Jianguo Lu and Baolong Bao.

## Abstract

Geoclimatic oscillations have repeatedly reshaped biodiversity, yet how ancient lineages persist and diversify through profound environmental upheavals remains a central question in evolutionary genomics. By integrating chromosome-level genome assemblies, population genomics, and transcriptomic data across the *Coilia* species complex, we reconstruct deep evolutionary histories of *Coilia* along the dynamic East Asian margins. Our analyses reveal divergences dating to the Miocene, coinciding with major tectonic and climatic transitions, including Tibetan Plateau uplift-linked reorganization of the Yangtze drainage toward the East China Sea, and the Middle Miocene Climatic Optimum. Despite long-term separation, we uncover evidence for ancient, reciprocal introgression among *Coilia* lineages, with gene flow confined to discrete genomic regions rather than distributed genome-wide. These introgressed regions are enriched for loci associated with immune function, vascular development, and osmoregulatory processes, and exhibit population genetic signatures consistent with positive selection. Temporal modelling indicates that this window of adaptive exchange preceded Pleistocene glacial intensification, after which gene flow became effectively restricted. Together, our results suggest that transiently permeable species boundaries historically acted as reservoirs of adaptive variation, shaping evolutionary trajectories and facilitating niche differentiation under repeated geoclimatic oscillations. This study highlights ancient adaptive introgression as an important evolutionary process contributing to the long-term persistence of threatened coastal fishes.

---

Geoclimatic oscillations have long been recognized as fundamental drivers of global biodiversity, particularly in dynamic landscapes where physical barriers periodically emerge and dissolve^1,2^. Classical evolutionary theory has traditionally emphasized allopatric isolation as the primary mechanism of lineage divergence^3,4^. However, accumulating genomic evidence increasingly demonstrates that speciation often proceeds through complex, reticulate processes rather than strictly bifurcating trajectories^5–7^. Within this framework, adaptive introgression has emerged as an important evolutionary process through which lineages may exchange genetic variation across partially permeable boundaries, potentially influencing ecological differentiation under shifting environmental conditions^8–11^. In aquatic systems characterized by strong environmental gradients, such genomic exchange has been repeatedly associated with enhanced evolutionary persistence during periods of profound geoclimatic change^12–15^. Nevertheless, the extent to which ancient introgression contributes to long-term adaptive potential across deep evolutionary timescales remains incompletely understood.

The East Asian marginal seas and their associated river systems constitute one of the most geologically dynamic regions on Earth, having been shaped by major tectonic events and repeated Pleistocene glacial cycles^16–18^. These processes generated steep and fluctuating environmental gradients, imposing substantial physiological and ecological constraints on aquatic organisms^19,20^. This region harbors the genus *Coilia*, a group of migratory fishes exhibiting pronounced ecological diversity, with species occupying freshwater, estuarine, and marine habitats^21,22^. Phylogenetic evidence indicates that *Coilia* lineages diverged tens of millions of years ago^22^, yet they persisted through dramatic environmental transformations, including the uplift of the Tibetan Plateau^23^ and the reorganization of regional ocean circulation patterns^24,25^. The combination of deep divergence, ecological heterogeneity, and inferred historical connectivity makes *Coilia* an informative system for investigating how geologically driven isolation and episodic genetic exchange jointly shape evolutionary trajectories across environmental boundaries.

Despite their evolutionary significance, populations of *Coilia*, such as the anadromous *C. nasus* and estuarine *C. mystus*, currently face severe declines due to overexploitation and habitat degradation, leading to their classification as threatened species^26,27^. Although phylogenetic relationships within the genus have been described^28^, the genomic processes underlying adaptation to heterogeneous and shifting environments remain poorly resolved. In particular, it is unclear how deeply diverged lineages overcame physiological and biotic challenges associated with salinity gradients, thermal variation, and pathogen exposure to occupy distinct ecological niches. Elucidating these historical adaptive processes is essential not only for understanding the evolutionary history of *Coilia*, but also for assessing the resilience of threatened aquatic species under ongoing environmental change.

Here, we generate a high-quality chromosome-level genome assembly for *C. mystus* and integrate population genomic, transcriptomic, and ecological niche modeling approaches to reconstruct the evolutionary history of the *Coilia* complex. Our analyses indicate that lineage divergence within *Coilia* coincided with major geoclimatic transitions in East Asia. We further identify genomic signatures consistent with ancient, reciprocal introgression among lineages, with introgressed regions concentrated in discrete portions of the genome rather than distributed genome-wide. These regions are enriched for loci associated with physiological processes relevant to immunity, vascular development, and osmoregulation, and exhibit population genetic patterns indicative of positive selection. By combining genomic inference with ecological niche modeling based on extensive occurrence data, we show that historical introgression coincided with shifts in habitat occupancy and environmental tolerance. Together, our results suggest that transiently permeable species boundaries historically acted as reservoirs of adaptive variation, shaping the evolutionary trajectories and likely contributing to the long-term persistence of threatened *Coilia* lineages under repeated geoclimatic oscillations.

## Results

### Distinct niche partitioning and environmental adaptations among *Coilia* species

The three *Coilia* species show marked spatial structuring along the East Asian coast (Fig. 1A). MaxEnt models recovered distinct core suitability areas for each species (Fig. 1B-D). *C. mystus* was predicted to occur broadly along the coast in nearshore habitats from tropical to temperate latitudes, while *C. nasus* and *C. grayii* formed a parapatric pair with a sharp north-south replacement. Suitability dominance profiling identified 25.60°N as the geographic transition for this parapatric pair (Fig. 1G): north of this latitude, predicted suitability was consistently higher for *C. nasus*, whereas *C. grayii* dominated in subtropical and tropical waters to the south.

**Figure 1.**
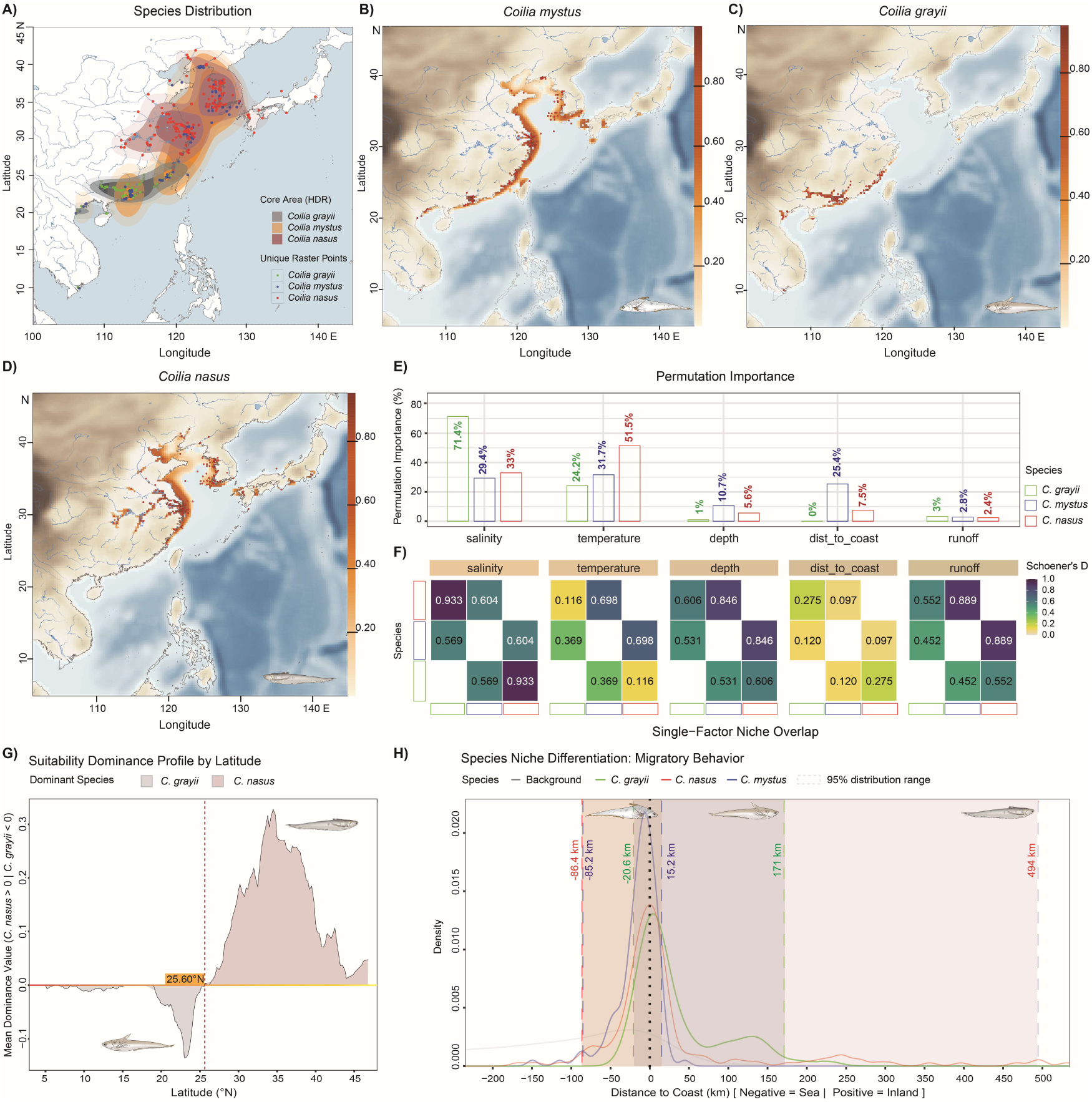
Geographic distribution, environmental drivers, and multidimensional niche differentiation in *Coilia*. **A)** Occurrence points and core distribution areas (50%, 75%, and 90% HDRs) estimated via non-parametric KDE. **B-D)** Predicted habitat suitability maps for *C. mystus*, *C. grayii*, and *C. nasus* generated by MaxEnt. **E)** Permutation importance of environmental variables for each species. **F)** Single-factor niche overlap (Schoener’s *D*) heatmaps. Statistical significance for niche equivalency (*p_Eq_*) and similarity (*p_Sim_*) was determined via 1,000 permutations; corresponding *p*-values are detailed in Supplementary Table S4. **G)** Latitudinal suitability dominance profile. The red dashed line indicates the geographic turnover threshold at 25.60°N. **H)** Kernel density distribution of signed distance to the coast. Shaded areas represent the 95% distribution intervals (95% CI), illustrating the distinct freshwater to marine migratory strategies across the three congeners.

Permutation importance highlighted different environmental associations among species (Fig. 1E). Salinity was the dominant predictor for *C. grayii* (71.4%), temperature for *C. nasus* (51.5%), and distance to the coast for *C. mystus* (25.4%), consistent with estuarine specialization in *C. grayii*, higher-latitude occupancy in *C. nasus*, and nearshore affinity in *C. mystus*.

Single-factor niche overlap analyses (Schoener’s *D*) further separated shared constraints from axes of divergence (Fig. 1F; Supplementary Table S4). Across all species pairs, salinity and depth niches were more similar than expected given the background (*p_Sim_* < 0.05; Supplementary Table S4), indicating a shared ecological baseline. In particular, salinity overlap between *C. grayii* and *C. nasus* was high (*D* = 0.933) and did not deviate from niche equivalency (*p_Eq_* = 0.1738; Supplementary Table S4). By contrast, overlap in surface runoff was moderate to high (e.g., *D* = 0.889 for *C. nasus* vs. *C. mystus*) but not more similar than expected by chance (*p_Sim_* > 0.05; Supplementary Table S4), suggesting that shared use of runoff-influenced habitats may primarily reflect environmental availability.

In contrast, temperature and signed distance to the coast showed pronounced divergence. Temperature comparisons yielded very low overlap for the parapatric pair (*C. grayii* vs. *C. nasus*, *D* = 0.116; Supplementary Table S4) and consistently rejected niche equivalency (*p_Eq_* = 1.000 across temperature pairwise tests; Supplementary Table S4). Signed distance-to-coast distributions further captured distinct habitat use consistent with contrasting migratory strategies (Fig. 1H). *C. nasus* spanned a broad freshwater-marine continuum (−86.4 km to 494 km), *C. grayii* occupied a narrow estuarine-riverine range (−20.6 km to 171 km), and *C. mystus* was concentrated near the estuarine-offshore interface (−85.2 km to 15.2 km). Correspondingly, distance-to-coast overlap was minimal between *C. nasus* and *C. mystus* (*D* = 0.097; *p_Eq_* = 1.000; Supplementary Table S4).

### A chromosome-level genome assembly reveals high chromosomal stability

We generated a chromosome-level reference genome for *C. mystus* (assembly size 893 Mb; longest scaffold 45.1 Mb; scaffold N50 34.8 Mb; Fig. 2A, B). In total, 94.8% of assembled sequences were anchored to 24 pseudochromosomes (Fig. 2A), providing a contiguous framework for chromosome-scale comparative analyses. Genome completeness was high based on BUSCO (*Actinopterygii_odb10*, n = 3,640; 95% complete, including 93.5% single-copy and 1.5% duplicated; 1.8% fragmented; 3.2% missing; Fig. 2B). K-mer-based profiling further supported a diploid genome with low residual heterozygosity and a modest sequencing error signal (Fig. 2C).

**Figure 2.**
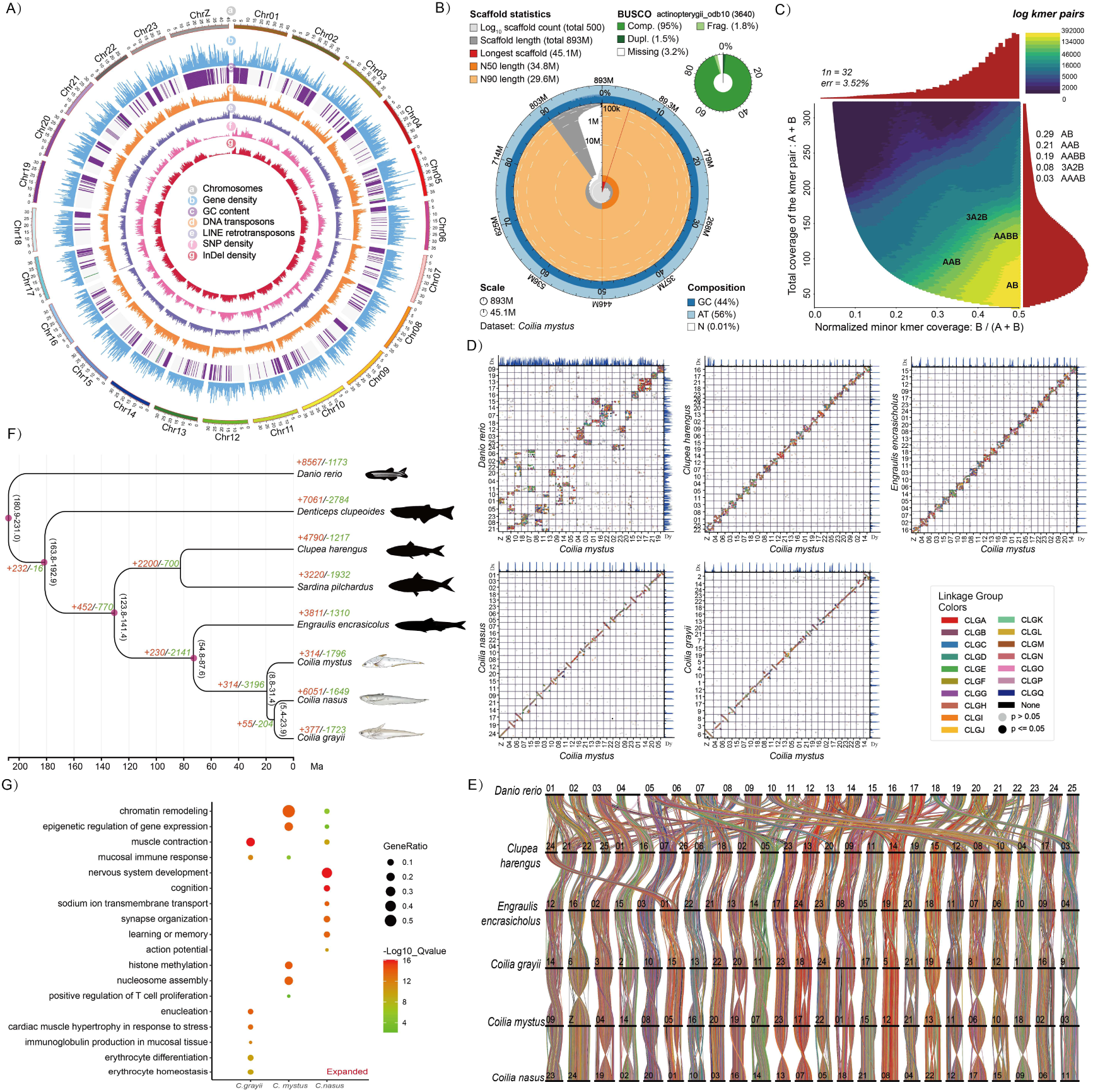
Chromosome-level genome assembly and comparative genomic landscape of *Coilia* species. **A)** Circos plot illustrating the genomic features of the *C. mystus* assembly across 24 pseudochromosomes. Rings from the outer to inner represent: (a) chromosomes, (b) gene density, (c) GC content, (d) DNA transposons, (e) LINE retrotransposons, (f) SNP density, and (g) InDel density. **B)** Assembly quality assessment showing scaffold length distribution and BUSCO completeness. **C)** Smudgeplot analysis of *k*-mer frequencies confirming the diploid state. **D)** Pairwise genome-wide synteny dot plots reconstructed using the protein-homology-based Odp suite, comparing *C. mystus* with *Danio rerio*, *Clupea harengus*, *Engraulis encrasicolus*, *C. nasus*, and *C. grayii*. Dots represent reciprocal best protein hits; significant syntenic blocks are colored. **E)** Macrosyntenic conservation across Clupeiformes visualized via ODP-inferred collinear ribbons. The diagram connects conserved syntenic blocks among the six analyzed teleost genomes, highlighting fusion and fission events during the evolution of the *Coilia* lineage. **F)** Chronogram reconstructed from single-copy orthologous genes. Purple nodes represent fossil-calibrated nodes used for molecular clock dating. Numbers at nodes indicate estimated divergence times (Ma); branch values (red/green) denote expanded and contracted gene families. **G)** Functional enrichment of expanded gene families. Dot size represents the GeneRatio, and color indicates the statistical significance (-log_10_*Q-value*).

Using single-copy orthologs, phylogenomic reconstruction placed *C. mystus* as sister to the (*C. grayii* + *C. nasus*) clade (Fig. 2F). Divergence time estimates dated the split between *C. mystus* and the other two *Coilia* lineages to the Miocene, followed by divergence between *C. grayii* and *C. nasus* (Fig. 2F). Despite pronounced ecological differentiation among species (Fig. 1), macro-synteny analyses showed largely one-to-one chromosome correspondences among the three *Coilia* genomes (Fig. 2D) and broadly conserved linkage relationships across representative Clupeiformes and an outgroup teleost (Fig. 2E), indicating limited large-scale chromosomal reshuffling over this timescale.

### Genomic signatures correlating with divergent niche adaptation

Comparative analyses of gene family expansions identified functional enrichments that mirror the ecological and migratory divergences among the three species (Fig. 2G; Supplementary Data 1). All three congeners shared expansions in categories related to ion transmembrane transport and muscle contraction, reflecting common physiological demands associated with estuarine-freshwater transitions.

In the long-distance migrant *C. nasus*, expanded families were enriched for neurobiological processes linked to spatial orientation and cognitive processing (e.g., learning, memory, and synapse organization; Fig. 2G). Enrichments also prominently featured muscle structural components and oxidative metabolism, paralleling the extensive energetic requirements of its migratory life history (Fig. 1H).

In *C. grayii*, expansions were concentrated in hematological and cardiopulmonary pathways (e.g., erythrocyte differentiation, cardiac muscle hypertrophy, elastic fiber formation; Fig. 2G). These genomic signatures are consistent with physiological adaptation to oxygen-limited, warm estuarine environments (Fig. 1C). Immune-related enrichments were also distinct, favoring categories associated with mucosal and barrier defenses.

In *C. mystus*, expansions were remarkably prominent in chromatin and epigenetic regulatory categories (e.g., chromatin remodeling, histone methylation, nucleosome organization; Fig. 2G). Additionally, immune-related enrichments involved systemic adaptive immunity (e.g., T-cell and NK cell proliferation). These expansions in regulatory and immune plasticity may underpin its capacity to maintain a broad coastal distribution across a wide latitudinal gradient (Fig. 1B).

### Population structure and genetic divergence among *Coilia* species

We analyzed 32,685,560 high-quality biallelic SNPs from 23 autosomes in 141 individuals spanning coastal and estuarine systems of East Asia (Fig. 3A). Both PCA and ADMIXTURE resolved three well-separated genetic clusters corresponding to *C. grayii* (CGR), *C. nasus* (CNA), and *C. mystus* (CMY) (Fig. 3B, C). PC1 (52.86%) primarily separated CMY from the CNA-CGR clade, whereas PC2 (37.38%) further distinguished CNA and CGR (Fig. 3C). Consistent with these patterns, the maximum-likelihood phylogeny also supported three monophyletic lineages matching species assignments (Fig. 3B).

**Figure 3.**
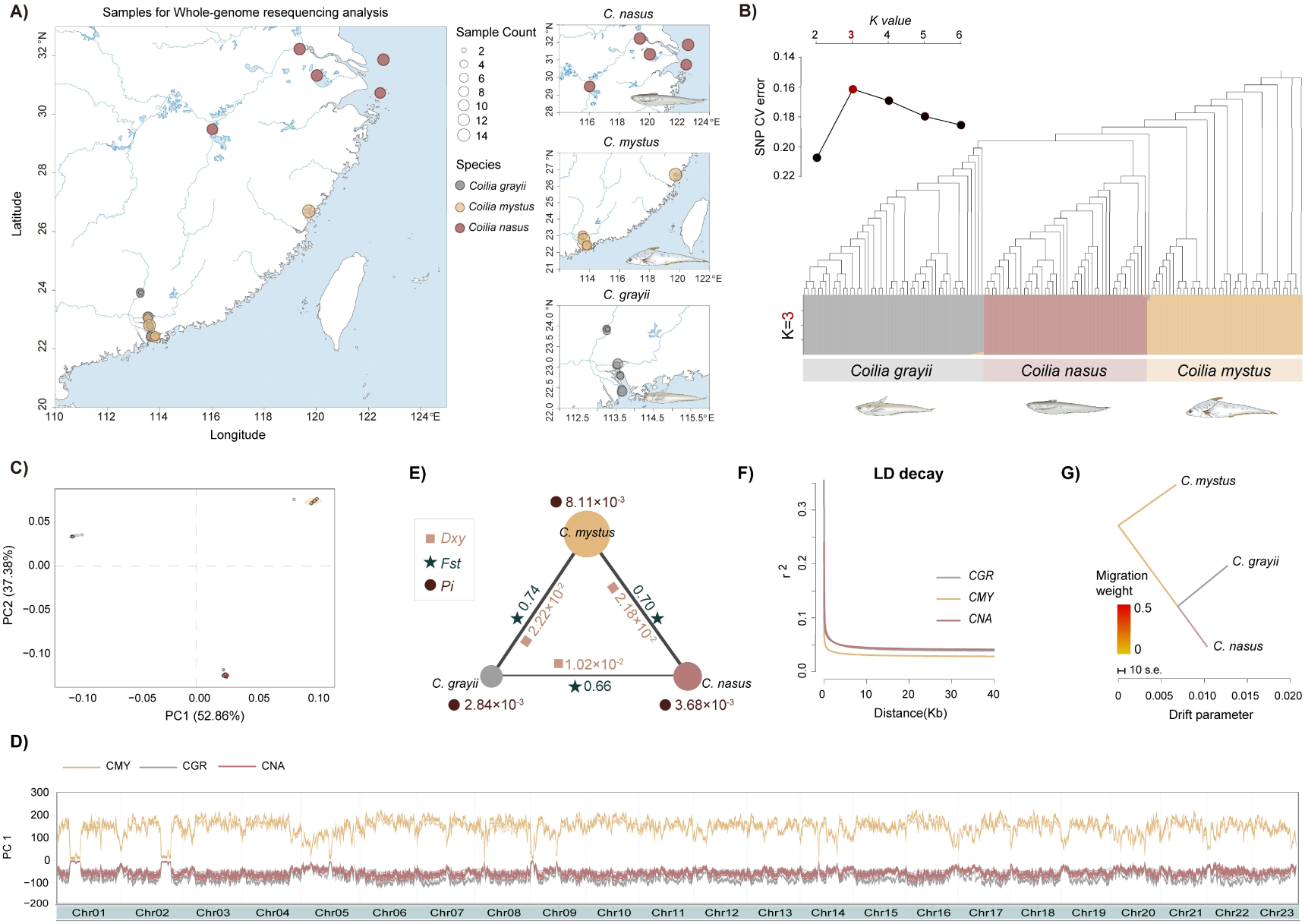
Population genomic architecture and lineage segregation in the genus *Coilia*. Analyses are based on 32,685,560 high-quality biallelic SNPs identified across 23 autosomes. **A)** Geographic sampling locations of the 141 individuals (51 *C. grayii*, 44 *C. mystus*, and 46 *C. nasus*) along the East Asian coastal and estuarine systems. **B)** Ancestry proportions inferred by ADMIXTURE at *K*=3 (bottom) and a maximum-likelihood phylogeny (top) for the 141-individual cohort. **C)** Principal component analysis (PCA) of the 141 individuals; the first two components (PC1 and PC2) explain 52.86% and 37.38% of the total genetic variance, respectively. **D)** Genome-wide PC1 scores calculated in 500-kb sliding windows with a step size of 10 kb across 23 pseudochromosomes, based on the refined dataset of 135 individuals with high lineage integrity. **E)** Triangle plot summarizing nucleotide diversity (pi, circles), genetic differentiation (*F_st_*, stars), and absolute divergence (*D_xy_*, squares) among species pairs (n = 135). **F)** Linkage disequilibrium (LD) decay curves showing the squared correlation of allele frequencies (*r*^2^) against physical distance (kb) for the three species. **G)** TreeMix inference based on the 135-individual dataset. Allowing 0-5 migration edges (m = 0-5) did not yield supported migration edges, consistent with limited detectable recent admixture among the three lineages.

To reduce potential bias in downstream demographic inference, we excluded six individuals showing minor ancestry components at *K* = 3 (maximum ancestry coefficient *Q* < 0.99), yielding a refined dataset of 135 individuals (Fig. 3B). Sliding-window PC1 values computed along the genome (500-kb windows, 10-kb step) remained stable across all 23 pseudochromosomes (Fig. 3D), indicating pervasive genome-wide differentiation rather than localized clustering driven by a small number of regions.

Genome-wide diversity and divergence metrics further highlighted strong lineage separation (Fig. 3E). CMY showed the highest nucleotide diversity (π = 8.11×10^-3^), whereas CGR and CNA exhibited lower diversity (π = 2.84×10^-3^ and 3.68×10^-3^, respectively). Pairwise differentiation was consistently high (*F_ST_* = 0.66-0.74), accompanied by substantial absolute divergence (*D_XY_* = 1.02×10^-2^ to 2.22×10^-2^) (Fig. 3E). Linkage disequilibrium decayed rapidly in all three species, with CMY showing the fastest decline in *r*^2^ with physical distance (Fig. 3F), consistent with its elevated diversity.

Finally, TreeMix analyses on the refined dataset supported a bifurcating species tree and did not infer migration edges under the tested setting (m = 0; Fig. 3G), suggesting no detectable recent, genome-wide admixture among the three lineages. Together, these results establish strong contemporary lineage segregation and provide a baseline for subsequent tests of deep-time introgression.

### Geologic and climatic drivers of divergence and gene flow

To reconstruct a demographic history robust to method-specific uncertainty, we adopted a hierarchical modeling strategy that integrates complementary temporal information. We used the chromosome-wise 95% confidence intervals (CIs) of gene-flow cessation inferred from SMC++ jackknifing together with phylogenetic divergence-time estimates from MCMCTree as informative priors to constrain fastsimcoal2 simulations (Fig. 4A, B). Under these biologically grounded constraints, model comparison decisively supported an "Ancient Gene Flow with Stepwise Isolation" (AGF-SI) scenario over alternative hypotheses of strict isolation or continuous migration (Supplementary Table S5). This integrative framework enabled us to anchor *Coilia* lineage dynamics to major geologic and climatic transitions along the East Asian margin (Fig. 4H).

**Figure 4.**
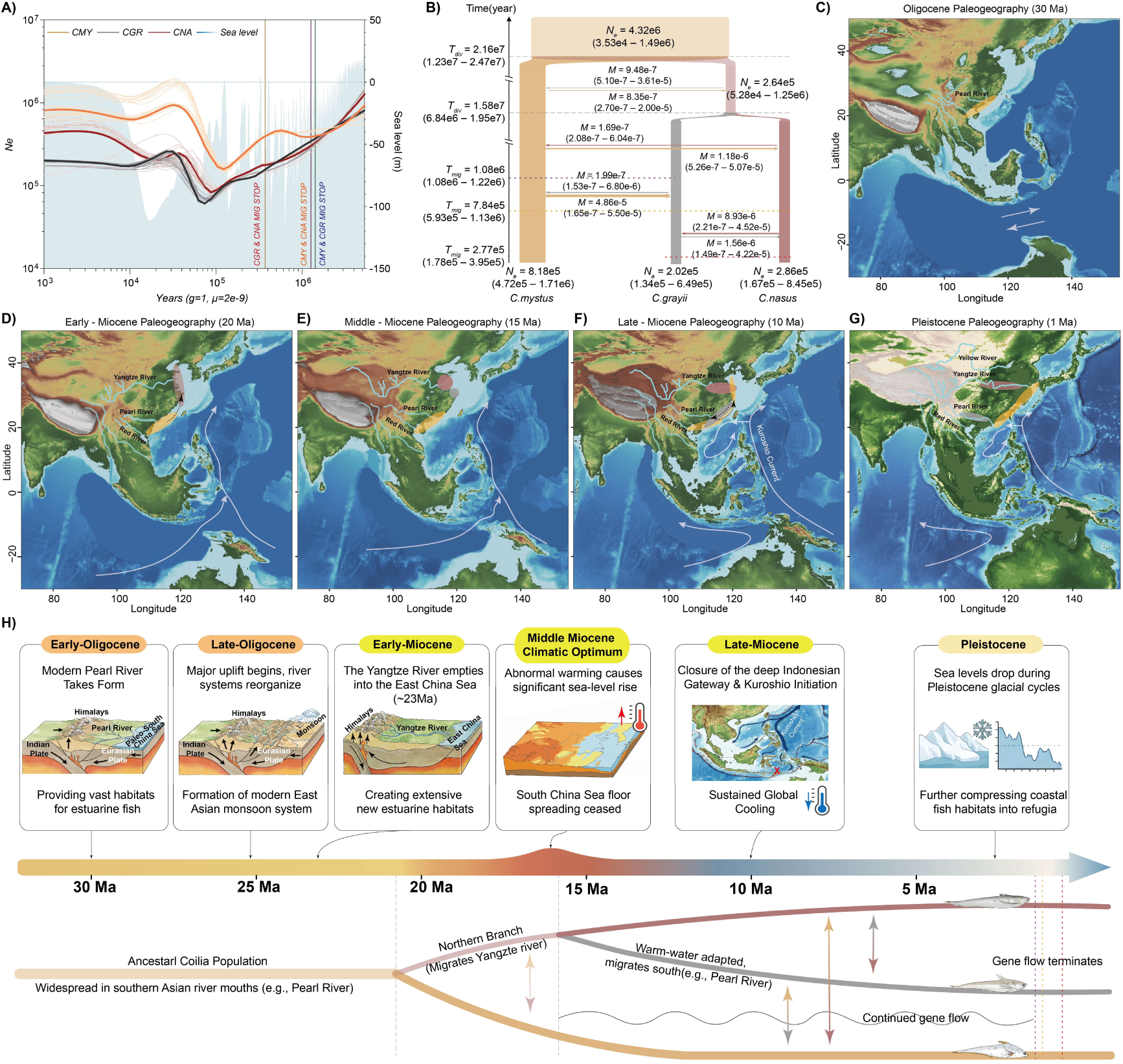
Demographic history and paleogeographic context of divergence and gene flow in *Coilia*. **A)** Effective population size (Ne) trajectories for *C. mystus* (CMY), *C. grayii* (CGR), and *C. nasus* (CNA) inferred using SMC++ over the last 3 million years. Thin background curves represent jackknife replicates, illustrating uncertainty in Ne estimates, and the blue shaded curve depicts global sea-level variation. Vertical colored lines indicate chromosome-wise estimates of gene-flow cessation times inferred independently across autosomes. **B)** Best-supported demographic scenario ("Ancient Gene Flow with Stepwise Isolation", AGF-SI) inferred using fastsimcoal2. The model depicts hierarchical divergence with asymmetric post-divergence gene flow and stepwise termination of exchange. Parameter labels show divergence times (Tdiv), migration rates (M), and cessation times (Tstop) with 95% CIs. Chromosome-wise cessation windows from A and divergence-time estimates from MCMCTree were used as informative priors to constrain the coalescent simulations. Arrow thickness reflects relative migration intensity. **C-G)** Paleogeographic reconstructions of the East Asian margin at 30 Ma **(C)**, 20 Ma **(D)**, 15 Ma **(E)**, 10 Ma **(F)**, and 1 Ma **(G)**. Dashed black arrows indicate hypothesized lineage dispersal routes associated with tectonic reorganization and climatic/oceanographic shifts; white arrows indicate major currents (e.g., Kuroshio) where shown. **H)** Schematic synthesis linking major geologic and climatic transitions (top) to inferred evolutionary milestones in *Coilia* (bottom), illustrating how tectonic reorganization and sea-level oscillations jointly shaped divergence, asymmetric gene flow, and its eventual termination.

Our paleogeographic reconstructions suggest that during the Oligocene (30 Ma), the ancestral Coilia population was likely confined to estuarine systems along the northern South China Sea margin (e.g., the paleo-Pearl River region)^29^, before the Yangtze River developed a through-flowing outlet to the East China Sea (Fig. 4C). Tectonic reorganization associated with uplift of the Tibetan Plateau likely promoted river integration and the establishment of the Yangtze drainage to the East China Sea around ∼23 Ma^30^, opening extensive new estuarine corridors. Coincident with this reconfiguration, the common ancestor of *C. grayii* and *C. nasus* expanded northward, initiating deep divergence from the *C. mystus* lineage at ∼21.6 Ma (95% CI: 12.3-24.7 Ma) (Fig. 4B, D).

Following this spatial expansion, the Middle Miocene (17∼14 Ma) was marked by two pivotal events: the Middle Miocene Climatic Optimum (MMCO), which caused sea levels to rise and submerge coastal plains (Fig. 4E)^31,32^, and the cessation of seafloor spreading in the South China Sea, which stabilized the basin morphology (Fig. 4E, H)^33^. These environmental shifts coincide with the estimated divergence between *C. grayii* and *C. nasus* at ∼15.8 Ma (95% CI: 7.2-20.9 Ma) (Fig. 4B). We propose this split was driven by ecological specialization: *C. nasus* adapted to long-distance anadromous migration into inland waters, while *C. grayii* retained a preference for warmer, estuarine environments. Following the MMCO, the subsequent Middle Miocene Climatic Transition (MMCT) led to an abrupt global cooling^34–36^. This climatic shift, coupled with the closure of the Indonesian Gateway (10 Ma)^24^, the consequent formation of the Western Pacific Warm Pool^37^, and the intensification of the East Asian Monsoon^38^, likely triggered the southward retreat of the warm-water adapted *C. grayii* and the concomitant expansion of *C. mystus* (Fig. 4F).

Despite these deep divergences, complex gene flow persisted until the Quaternary. Our demographic modeling quantified a marked asymmetry in genetic exchange (Fig. 4B). As the Pleistocene glaciations intensified (2.58 Ma), SMC++ analysis reveals a synchronous decline in effective population size (*Ne*) across all lineages (Fig. 4A), tightly mirroring the progressive sea-level regression that fragmented continuous estuarine habitats into isolated refugia (Fig. 4G)^39,40^. This habitat fragmentation ultimately led to the hierarchical cessation of gene flow. The striking contrast between the deep phylogenetic divergence inferred by MCMCTree and the recent cessation of genetic exchange revealed by SMC++ implies a prolonged history of semi-permeable species boundaries. While the overall magnitude of gene flow was restricted, preserving the specific identity of each lineage, it remained sufficient to facilitate the transfer of advantageous alleles. Crucially, these precise demographic parameters, particularly the divergence times and asymmetric migration rates, established a robust null model for our subsequent machine-learning-based detection of introgressed genomic windows (FILET).

### Asymmetric and functionally biased adaptive introgression

To characterize the genomic consequences of ancient gene flow during *Coilia* divergence, we applied FILET using simulations parameterized by the demographic history inferred above. This scan identified multiple high-confidence introgressed tracts across the autosomes (posterior probability ≥ 0.99; Fig. 5A), with strongly asymmetric directionality. *C. mystus* contributed the largest fraction of introgressed sequence, with 25.2 Mb (3.12%) assigned to *C. grayii* and 9.8 Mb (1.21%) to *C. nasus*. In contrast, introgression into *C. mystus* was comparatively limited from *C. grayii* (4.3 Mb; 0.53%) and minimal from *C. nasus* (0.4 Mb; 0.048%). To highlight candidate adaptive events, we intersected introgressed tracts with the upper tail of haplotype-based selection statistics (top 5% iHS and nSL), yielding a set of high-confidence loci in which introgression and selection signatures co-localize (Fig. 5B).

**Figure 5.**
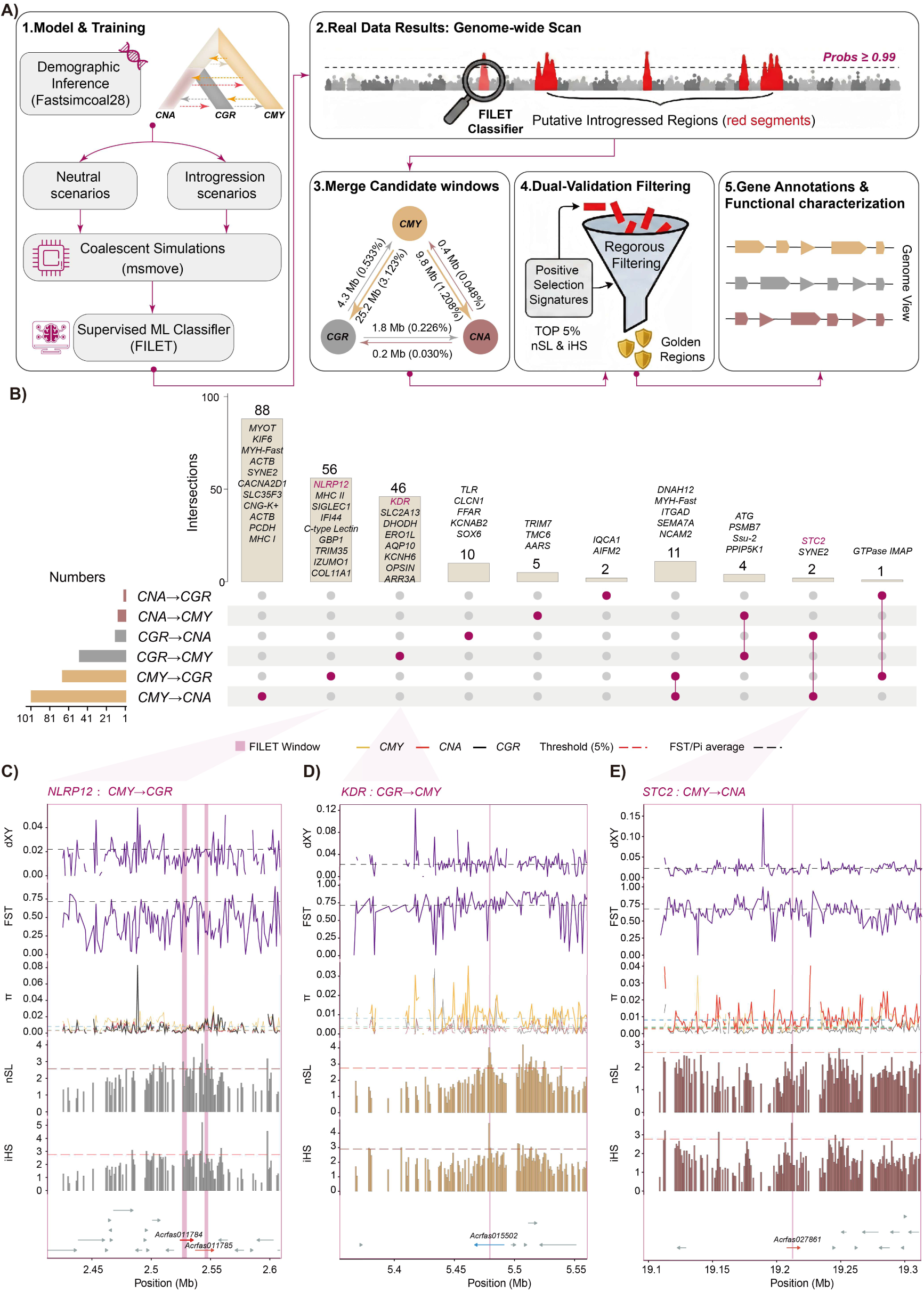
Genome-wide inference and prioritization of candidate adaptive introgression in the *Coilia* complex using a supervised machine-learning framework. **A)** Workflow for detecting introgressed tracts. Demographic parameters inferred with fastsimcoal2 were used to simulate neutral and introgression scenarios (msmove) for training the supervised classifier FILET. The trained classifier was then applied to real genomes to identify putative introgressed regions (posterior probability ≥ 0.99; red segments), which were merged into candidate windows. Candidate windows were further prioritized by intersection with haplotype-based selection signals (top 5% nSL and iHS), yielding high-confidence candidate "Golden Regions", followed by gene annotation and functional characterization. The inset triangle summarizes the total genomic length and proportion of inferred introgressed tracts between donor-recipient directions. **B)** UpSet plot showing genes overlapped by inferred introgressed windows across donor-recipient directions. Bars indicate the number of genes in each set or intersection; the matrix denotes the corresponding directionality. Key candidates highlighted in the text (e.g., *NLRP12*, *KDR*, *STC2*) are annotated. **C-E)** Local genomic landscapes of three representative candidate loci: *NLRP12* (CMY→CGR; **C**), *KDR* (CGR→CMY; **D**), and *STC2* (CMY→CNA; **E**). Tracks show (top to bottom) *D_xy_*, *F_ST_*, π, and haplotype-based statistics (nSL and iHS). Pink shading indicates FILET-inferred introgressed windows. Dashed lines denote genome-wide reference levels and the top 5% thresholds for nSL/iHS. Note that *STC2* is detected in introgression into CNA from both estuarine lineages (CMY and CGR; see panel B), while panel E shows the CMY→CNA window as a representative example.

Introgression from *C. mystus* into *C. grayii* was enriched for immune- and defense-associated candidates, including loci in the MHC and TRIM families (Supplementary Table S6&S7). A representative example, *NLRP12*, falls within an introgressed tract and shows elevated haplotype statistics (Fig. 5C), consistent with selection on immune-related variation in estuarine environments. By contrast, introgression into the anadromous *C. nasus* highlighted candidates linked to osmoregulatory and ion-transport processes, including ion-handling genes (*CACNA2D1*, *MICU3*) and solute carriers (Supplementary Table S6&S7). The strongest co-localized signal involved *STC2*, with introgressed tracts inferred from both estuarine lineages (*C. mystus* and *C. grayii*), suggesting reuse of a shared estuarine-associated variant (Supplementary Table S7). At this locus, reduced nucleotide diversity (π) together with elevated iHS/nSL supports a pronounced sweep-like pattern in *C. nasus* (Fig. 5E).

In the reverse direction (*C. grayii* to *C. mystus*), we detected introgressed tracts overlapping developmental candidates such as *KDR* (VEGFR2) and *SNTB1*. (Supplementary Table S7). The *KDR* region similarly exhibits co-localized haplotype-based signatures (Fig. 5D), consistent with selection acting on introgressed variation potentially relevant to vascular or hypoxia-related physiology. Collectively, these results indicate that, despite deep divergence, historical gene flow left a quantitatively asymmetric footprint and that a subset of introgressed tracts co-localize with strong haplotype-based signals at loci plausibly related to ecological differentiation across the *Coilia* complex.

### Transcriptomic reshaping and regulatory divergence at introgressed loci

To evaluate whether candidate introgressed loci show regulatory divergence across contrasting hydrological regimes, we profiled muscle transcriptomes from four wild groups sampled along a salinity and hydrodynamic gradient: freshwater (Yangtze River; *C. nasus*), semi-enclosed and relatively stable coastal waters (Sansha Bay; North *C. mystus*), and dynamic estuarine environments (Pearl River Estuary; South *C. mystus* and *C. grayii*) (Fig. 6A). PCA separated samples primarily by habitat along PC1 (39% variance; freshwater vs. estuarine), whereas PC2 (24%) captured within-*C. mystus* structure consistent with the north-south contrast in hydrodynamic regime (Fig. 6A).

**Figure 6.**
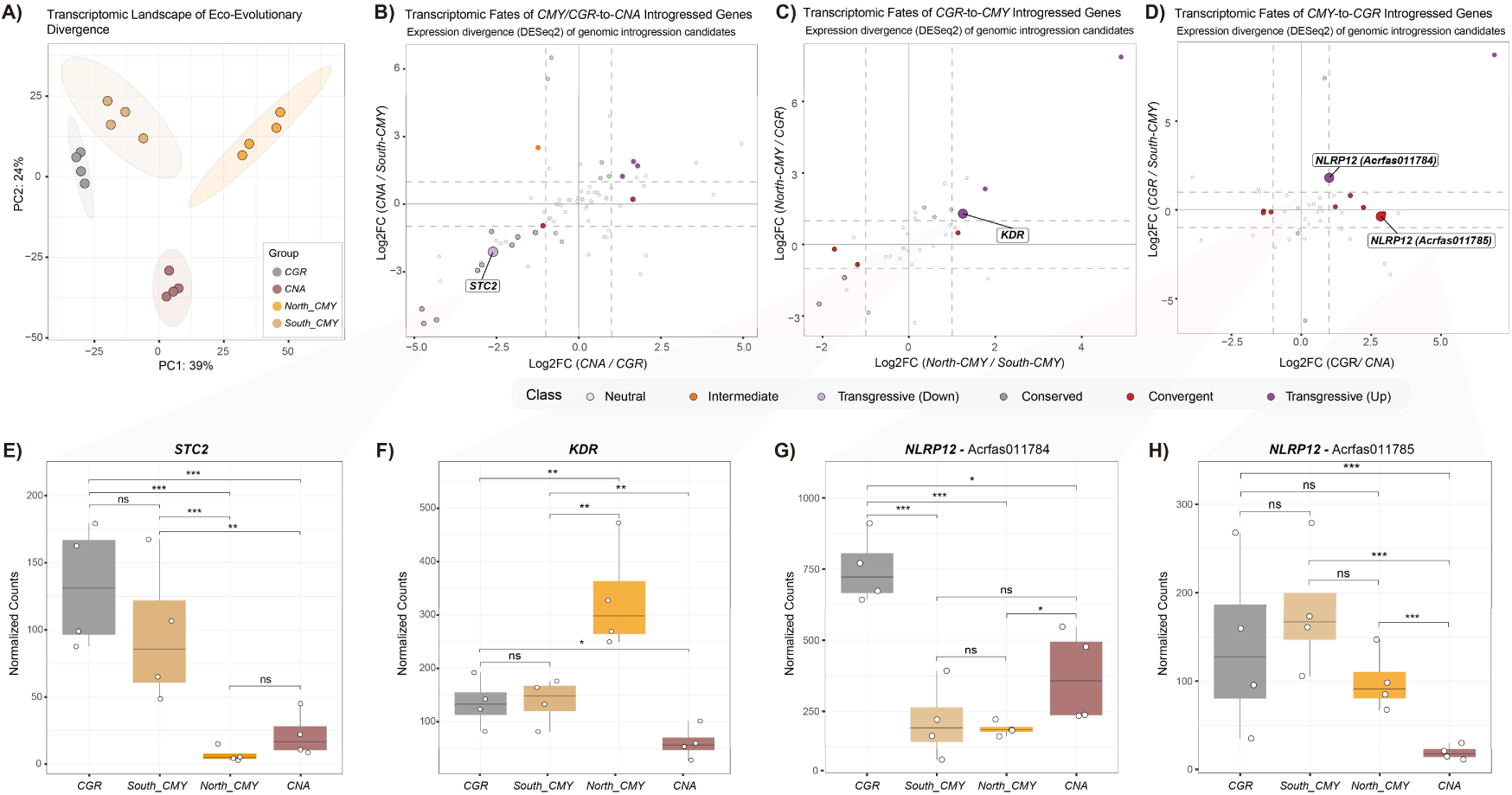
Transcriptomic landscape and regulatory divergence of introgressed loci across hydrological regimes. **A)** PCA of muscle transcriptomes from *C. nasus* (CNA), *C. mystus* (North_CMY and South_CMY), and *C. grayii* (CGR). PC1 (39%) separates freshwater CNA from estuarine lineages, and PC2 (24%) captures within-*C. mystus* structure. **B-D)** regulatory fates of candidate genes located in introgressed windows. Scatter plots compare expression divergence (DESeq2 log2FC) in the putative recipient lineage relative to the putative donor (x-axis) and to the non-introgressed sister lineage (y-axis). Points are categorized as conserved, convergent, or transgressive based on divergence relative to both comparators. Dashed lines indicate |log2FC| = 1 thresholds. B) CNA comparison highlighting *STC2*. C) North_CMY comparison highlighting *KDR*. D) CGR comparison highlighting *NLRP12* paralogs (Acrfas011784 and Acrfas011785). **E-H)** Normalized expression counts (DESeq2) for key adaptive candidates. **E)** *STC2* maintains high expression in populations inhabiting dynamic estuarine environments (CGR and South_CMY) but is significantly downregulated in freshwater CNA and the stable, semi-enclosed North_CMY. **F)** *KDR* exhibits significant upregulation specifically in the high-latitude North_CMY population. **(G, H)** *NLRP12* paralogs display distinct expression patterns in CGR, indicative of sub-functionalization. Box plots represent the median (center line), upper and lower quartiles (box limits), and 1.5× interquartile range (whiskers). Statistical significance was determined using pairwise Wald tests, with *P*-values adjusted for multiple testing (Benjamini-Hochberg; * *P*_adj_ < 0.05, ** *P*_adj_ < 0.01, *** *P*_adj_ < 0.001; ns, not significant).

Focusing on genes located within introgressed windows, we compared expression divergence in putative recipient lineages relative to donor and non-introgressed sister lineages and classified regulatory outcomes as conserved, convergent, or transgressive (Fig. 6B-D). Across loci, introgressed candidates were frequently associated with non-conserved expression trajectories, indicating that introgressed haplotypes can be accompanied by lineage- and habitat-dependent regulatory shifts.

At the osmoregulatory candidate *STC2*, expression in *C. nasus* was significantly transgressively downregulated relative to both estuarine lineages (Fig. 6B, E; *P_adj_* < 0.001). Within *C. mystus*, *STC2* also differed between populations, with higher expression in the estuarine South_CMY and lower expression in the more stable North_CMY (Fig. 6E), consistent with spatially structured regulation across contrasting hydrological contexts.

For the vascular-development candidate *KDR*, the North_CMY population exhibited significant transgressive upregulation (Fig. 6C; *P_adj_* < 0.001), with expression exceeding both South_CMY and the putative donor lineage (Fig. 6F; pairwise *P_adj_* < 0.01). In addition, the immune-associated locus *NLRP12* displayed divergent regulatory fates between paralogs in *C. grayii*: one copy (Acrfas011785) showed a convergent pattern, whereas the other (Acrfas011784) showed transgressive upregulation (Fig. 6D, G, H), consistent with regulatory divergence following introgression.

## Discussion

Our genomic reconstruction shows that the evolutionary trajectory of the *Coilia* complex has been fundamentally orchestrated by the interplay of tectonics and climatic oscillations in East Asia. In contrast to the rapid, recent radiations characteristic of many post-glacial freshwater fishes, *Coilia* lineages exhibit deep divergence times dating back to the Miocene. This chronology aligns precisely with major geological events, specifically the reorganization of East Asian drainage systems linked to the uplift of the Tibetan Plateau^42–44^. The initial split between the *C. mystus* lineage and the ancestor of *C. grayii* and *C. nasus* coincides with the establishment of the modern Yangtze River drainage^30^, which likely provided novel freshwater conduits and estuarine habitat facilitating initial allopatric speciation.

While tectonic events set the stage, subsequent climatic oscillations dictated the specific ecological trajectories of these lineages. We posit that the Middle Miocene Climatic Optimum (MMCO, ∼15 Ma) established a warm thermal baseline that drove the ecological differentiation of the *C. grayii*-*C. nasus* ancestor^32,44,45^. However, the biological landscape was drastically remodeled by the subsequent Middle Miocene Climate Transition (MMCT), characterized by rapid global cooling and the intensification of the East Asian Monsoon. Concurrently, the concurrent progressive constriction of the Indonesian Gateway (reaching a critical threshold ∼10 Ma) restricted the Indonesian Throughflow, fostering the formation of the Western Pacific Warm Pool and its thermal extension into the South China Sea^37,46,47^. This restructured oceanic thermal gradient forced the warm-adapted *C. grayii* to retreat southward into the stable thermal refugium of the subtropical Pearl River basin. This geoclimatically driven range shift set the stage for secondary contact, transforming the genus from a simple bifurcation model into a complex reticulate system.

These profound ecological shifts left distinct footprints on the functional architecture of each lineage, serving as genomic archives of their long-term specialization. For the anadromous *C. nasus*, the unique expansion of neurobiological and musculoskeletal pathways parallels the extensive migratory demands of the Yangtze River corridor^48,49^. These adaptations likely provided the physiological blueprint for conquering the energetic and cognitive demands of the longest riverine migration in East Asia. Conversely, the southern-restricted *C. grayii* exhibits expansions in hematopoietic and cardiac functions, consistent with physiological acclimatization to the warm, periodically hypoxic habitats of the Pearl River system^50,51^. We posit that these specialized traits served as a critical survival toolkit, enabling persistence in a region where thermal stress and hypoxia act as potent ecological filters. Finally, the basal *C. mystus* shows a disproportionate expansion of epigenetic regulatory machinery, reflecting a strategy of phenotypic plasticity to cope with the extreme environmental heterogeneity of the coastline^52,53^. This regulatory flexibility likely constitutes a fundamental mechanism supporting its exceptional capacity to span the entire latitudinal gradient of the East Asian coast, effectively buffering against diverse physiological stressors.

Within this established ecological framework, a central finding of our study is that ancient adaptive introgression may have contributed to lineage persistence by enabling bidirectional exchange of functionally relevant alleles during geoclimatic upheavals. We detect reciprocal ancient introgression concentrated in candidate loci with functions plausibly linked to adaptation across contrasting environments. In subtropical estuarine habitats, *C. grayii* likely encountered pathogen communities distinct from those in temperate coastal systems. In line with this ecological setting, our genomic analyses highlight introgression at immune-related candidates, including *NLRP12*^54,55^, with *C. mystus* as a putative source lineage. Although fitness effects cannot be quantified directly here, the known role of *NLRP12* in innate immune regulation^54,55^ provides a plausible explanation for why introgressed immune variants may have been favored under local microbial pressures, potentially accelerating adaptation relative to reliance on de novo mutation^56,57^.

Reciprocal signals in the opposite direction further indicate that introgression was not restricted to immune-related loci, but also involved candidates associated with physiological function. We infer reciprocal introgression at the vascular development locus *KDR* (VEGFR2), with *C. grayii* as a putative donor lineage and *C. mystus* as the recipient. Given the established role of *KDR* in angiogenesis and oxygen delivery^59–61^, this pattern is consistent with the hypothesis that introgressed variation could have contributed to physiological flexibility in vascular function. Notably, *KDR* shows elevated expression in northern *C. mystus* populations, providing supportive functional context for population-specific regulatory differentiation, although it does not by itself establish allele-specific effects. Together, these observations suggest that introgressed variation, followed by regulatory tuning, may have facilitated the expansion across a broad latitudinal distribution of *C. mystus* coinciding with the establishment of modern regional circulation patterns^47^. In this view, introgression could have supplied functionally relevant variants that were subsequently filtered during niche divergence, potentially supporting persistence of *C. grayii* in variable southern estuaries and broader niche breadth in *C. mystus*.

Beyond thermal and immune adaptations, our results also provide insight into the genomic correlates of anadromy in *C. nasus*. We detect introgression signals at *STC2*, with both estuarine lineages (*C. mystus* and *C. grayii*) representing plausible source lineages, consistent with incorporation of a haplotype associated with estuarine environments. *STC2* is a key regulator of calcium and phosphate homeostasis^61,62^, making it a biologically plausible candidate for osmotic transitions. However, introgression alone is unlikely to account for the anadromous phenotype. Instead, our transcriptomic data reveal a pronounced, recipient-lineage-specific regulatory shift:

*STC2* is significantly downregulated in freshwater *C. nasus* relative to both comparison lineages, yielding a transgressive expression pattern. This habitat-associated downregulation is consistent with regulatory modification of ion homeostasis pathways under hypo-osmotic conditions^63^, suggesting that introgressed genetic scaffolds can be repurposed through subsequent regulatory change. More broadly, this result supports the view that adaptive introgression may act in concert with regulatory innovation to generate phenotypes beyond the range observed in donor and sister lineages^64^, potentially facilitating the use of extreme environments such as freshwater migration corridors^65^.

Together, these candidate loci illustrate that introgression was not genome-wide, but instead occurred in a restricted and potentially functionally biased manner, prompting a broader view of species boundaries in *Coilia*. The apparent discordance between deep phylogenetic divergence and comparatively recent cessation of gene flow is consistent with *Coilia* lineages maintaining semi-permeable species boundaries over extended timescales. This prolonged "grey zone" of speciation challenges the classical view of reproductive isolation as a rapid and irreversible transition^66^. Rather than strict bifurcation, our results support a scenario in which related lineages remained partially connected, resembling elements of a syngameon-like system in which a species complex can act as a reservoir of standing variation^67^. In such a context, restricted yet persistent exchange may permit the episodic incorporation of potentially beneficial variants while limiting genome-wide homogenization, thereby preserving lineage integrity. Importantly, this permeability appears to have been curtailed during the Quaternary: intensified Pleistocene glaciations and sea-level regression likely fragmented estuarine habitats, promoting stepwise isolation and reducing interlineage gene flow. In this sense, Pleistocene habitat fragmentation may have locked in earlier introgressed variation as persistent genomic legacies within each lineage.

This reticulate history also provides a conservation-relevant context. The pronounced genomic differentiation among modern lineages suggests that contemporary anthropogenic pressures (e.g., barriers to migration, habitat degradation, and overexploitation) may be acting on populations whose evolutionary histories were shaped under periodically connected river-estuary-coastal landscapes^68^. The ancient introgressed alleles that once facilitated niche colonization are now fixed legacy traits. However, the complete cessation of gene flow in the late Pleistocene means that modern populations lack the potential for genetic rescue via natural hybridization^69,70^. Consequently, the rapid loss of genetic diversity in these threatened species is particularly alarming, as it erodes the finite standing variation derived from millions of years of evolutionary trial-and-error. Conservation strategies must therefore prioritize the maintenance of large effective population sizes and the protection of migratory corridors, treating these distinct genomic lineages as crucial management units to preserve their remaining adaptive potential.

In summary, our integrative genomic and ecological analyses suggest that the *Coilia* complex reflects deep geological divergence accompanied by episodic, reciprocal introgression across semi-permeable species boundaries. Introgression signals are concentrated in candidate loci with functions plausibly linked to salinity, thermal, and biotic challenges, providing a mechanistic context for niche differentiation under past geoclimatic instability. More broadly, our results highlight how long-lived lineages can retain partially reticulate histories and preserve adaptive variation despite deep divergence. As human activities increasingly reshape river-estuary-coastal connectivity and environmental regimes, placing contemporary vulnerability in a deep-time demographic framework can help frame expectations about adaptive capacity and inform conservation of threatened aquatic lineages.

## Methods

### Sample collection and study design

Biological sampling was conducted between September 2023 and September 2025 under approval of the Animal Care and Use Committee of Sun Yat-sen University. For de novo genome assembly, we collected a single male *C. mystus* to ensure uniform coverage of the Z chromosome under the reported ZZ/ZO sex determination system in this species^71^.

For population resequencing, muscle tissues preserved in 95% ethanol were obtained from representative habitats: *C. nasus* from Poyang Lake and the Zhoushan Archipelago, supplemented with publicly available genomic data from freshwater and marine populations (BioProject PRJNA744015)^72^; *C. mystus* from Sansha Bay and the Pearl River Estuary; and *C. grayii* from the Pearl River Estuary and the Beijiang River (Yingde section).

For transcriptomic analyses, we generated new RNA-seq data from flash-frozen muscle tissues of *C. mystus* (Sansha Bay) and *C. mystus*/*C. grayii* (Pearl River Estuary). Public muscle transcriptomes of *C. nasus* (BioProject PRJNA1273475)^73^ were retrieved from the NCBI Sequence Read Archive. Sampling locations (Fig. 3A), accession numbers, and metadata are provided in Supplementary Table S1.

### Species occurrence data and spatial thinning

Occurrence records for *C. nasus*, *C. grayii*, and *C. mystus* were compiled from field surveys, the Global Biodiversity Information Facility, and literature sources accessed via Google Scholar and the China National Knowledge Infrastructure.

Records were restricted to the period 1990-2025. To reduce spatial sampling bias, we applied spatial thinning, retaining one occurrence per 0.25° × 0.25° grid cell. Core distribution areas were estimated as Highest Density Regions using kernel density estimation implemented in the ggdensity package^74^.

### Environmental framework and signed distance logic

We constructed an environmental stack comprising five core variables. Temperature and surface runoff were extracted from ERA5 monthly averaged reanalysis data (1993-2025)^75,76^. Salinity data were obtained from GLORYS12V1 via the Copernicus Marine Service^77^; missing salinity values in riverine or estuarine cells (depth ≤ 0) were assigned a value of 0. Bathymetry was derived from the GEBCO 2025 grid^78^, with land cells containing confirmed occurrences adjusted to −1 m. A signed distance to the coast was calculated using a Natural Earth land mask, with positive values assigned to inland areas and negative values to marine areas. Species-specific ecological ranges were summarized using the 95% distribution intervals of signed-distance values.

### Species distribution modeling and niche analysis

Potential habitat suitability was modeled using the Maximum Entropy algorithm (MaxEnt v3.4.4)^79^. The accessible area was defined by a 5° buffer around occurrence locations. Variable importance was assessed using permutation importance, and binary suitability maps were generated using the 10th percentile training presence threshold^80^. Subsequently, to quantify the latitudinal replacement between *C. nasus* and *C. grayii*, we utilized continuous suitability outputs to generate latitudinal dominance profiles.

Model performance was evaluated using the area under the receiver operating characteristic curve (AUC). All three species models exhibited high predictive accuracy, with AUC values ranging from 0.961 to 0.990 (Supplementary Figure S1), indicating excellent discriminatory power and low overfitting risks.

Niche overlap and shifts were quantified using the Broennimann framework^81^ implemented in the ecospat package^82^. Occurrence densities were estimated in PCA-calibrated environmental space using kernel density estimation (R = 1000). Niche overlap was quantified using Schoener’s D^83^. Equivalency and similarity tests followed Rödder & Engler^84^ and were conducted using 1,000 permutations with a fixed random seed (1024).

### Genome sequencing, assembly and annotation

High-molecular-weight DNA from a male *C. mystus* specimen was sequenced on the PacBio Revio platform (CCS mode) to generate HiFi reads. Hi-C libraries were constructed from muscle tissue and sequenced on the MGISEQ-2000 platform. De novo assembly was performed using Hifiasm (v0.19.8)^85^, followed by haplotypic duplication removal with Purge_Dups (v1.2.5)^86^. Hi-C reads were mapped using BWA^87^ and scaffolding was conducted using Yahs (v1.1)^88^. Manual curation was performed in PretextView. Assembly completeness was assessed using BUSCO (v5.4.7)^89^ and GenomeScope (v2.0)^90^.

Sex chromosomes were identified by comparing normalized sequencing depth across 30 resequenced individuals using Mosdepth (v0.3.3)^91^. To correct for sequencing output variations, the mean depth of each chromosome was normalized against the average depth of all 23 autosomes within each individual (Supplementary Table S2). Repetitive elements were annotated using RepeatModeler (v2.0.5)^90^ and RepeatMasker (v4.1.6)^92^. Protein-coding gene prediction was performed using the GETA pipeline (v2.5.3), integrating ab initio prediction, homology-based evidence, and transcriptomic data. Homology-based evidence used protein sets from clupeiform genome assemblies, including *Sardina pilchardus* (GCF_963854185.1), *Engraulis encrasicolus* (GCF_034702125.1), *Alosa sapidissima* (GCF_018492685.1), and *C. nasus* (GCA_027475355.1). Transcriptomic evidence was derived from a pooled multi-tissue RNA-seq library and mapped using HISAT2^93^ within the GETA pipeline. Functional annotation was conducted using SwissProt, NR, Pfam, KEGG, and Gene Ontology databases via eggNOG-mapper^94^.

### Phylogenomics, divergence time estimation, and gene family dynamics

Phylogenomic analyses included genome assemblies from eight species: *Danio rerio* (GCF_000002035.6)^95^, *Denticeps clupeoides* (GCF_900700375.1)^96^, *Clupea harengus* (GCF_900700415.2)^97^, *Sardina pilchardus* (GCF_963854185.1)^98^, *Engraulis encrasicolus* (GCF_034702125.1), *C. nasus* (GCA_027475355.1)^99^, *C. grayii* (GCA_042479465.1)^100^, and *C. mystus* (this study). Single-copy orthologs were identified using OrthoFinder^101^. Protein alignments were generated with MUSCLE^102^, and codon alignments were produced using MACSE^103^. Species topology was inferred using the STAG algorithm.

Divergence times were estimated using MCMCTree (PAML v4.9)^104^ under an independent rates clock (clock = 2) using the LG substitution model¹⁰⁵ with gamma rate heterogeneity (model = 2, alpha = 0.5). We ran the MCMC for 50,000 samples after a burn-in of 3,000,000 iterations, sampling every 1,000 iterations. Fossil calibrations followed Egan et al.^105^ and were applied to four nodes: Root (Otocephala) 151-228 Ma; Clupeiformes 137-190 Ma; Crown Clupeoidei 125-145 Ma; Crown Engraulidae 50-86.3 Ma. Gene family expansions and contractions were inferred using CAFE5^106^ under a stochastic birth-death model. Input gene counts were derived from OrthoFinder orthogroups, with families present in fewer than two species excluded to ensure phylogenetic informativeness. The global birth-death parameter (λ) was estimated using maximum likelihood, incorporating an error model to account for potential assembly and annotation artifacts. Families exhibiting significant size changes were identified using a conditional P-value threshold of < 0.05 (Viterbi method). Functional enrichment analyses of these significantly expanded or contracted families were performed using GO enrichment implemented in TBtools-II^107^, applying a Benjamini-Hochberg corrected FDR < 0.05^108^.

### Synteny and chromosomal evolution analysis

Comparative synteny analyses were performed between *C. mystus* and five teleost genomes (*Danio rerio*, *Clupea harengus*, *Engraulis encrasicolus*, *C. nasus*, and *C. grayii*) using Oxford Dot Plots (ODP)^109^. Putative orthologs were identified via reciprocal best protein matches using Diamond BLASTP^110^. Pairwise genome-wide dot plots and multi-genome collinear ribbons were generated, and syntenic associations were assessed using Fisher’s exact test and the D statistic^111^.

### Read mapping, variant calling, and filtering

Genomic DNA was extracted from muscle tissues of 111 newly sampled individuals using a modified phenol-chloroform protocol. Libraries (200-400 bp inserts) were sequenced on the DNBSEQ-T7 platform as 150 bp paired-end reads with ∼10× mean coverage per individual. Public resequencing data for 30 *C. nasus* individuals were retrieved from BioProject PRJNA744015 and processed jointly. Raw reads were filtered using fastp^112^ (adapter trimming; Q < 20; minimum length 50 bp). Clean reads were mapped to the *C. mystus* reference using BWA-MEM^113^, duplicates were marked using Picard, and alignment statistics were summarized using Samtools^114^. Variants were called following GATK Best Practices using GATK (v4.5)^115^ HaplotypeCaller in GVCF mode, followed by joint genotyping (GenomicsDBImport; GenotypeGVCFs). Variants were filtered using hard quality thresholds, population-level depth and missingness criteria via VCFtools^116^, we retained biallelic SNPs with a minimum depth of 5, a maximum depth of 25, and a genotyping rate > 80% (--max-missing 0.8). (3) Mappability masking used GenMap (v1.3.0)^117^ to calculate genome mappability with k-mer size 150 bp allowing two mismatches; regions with mappability < 1 were merged using bedtools^118^, and variants in masked regions were removed using BCFtools^119^.

### Population structure and phylogenetic analysis

To reduce the impact of physical linkage on multivariate and clustering analyses, we generated an LD-pruned SNP set using PLINK2^120^ with --indep-pairwise 50 5 0.2 (window size 50 SNPs, step size 5 SNPs, r² ≥ 0.2), excluding non-autosomal contigs. PCA and ADMIXTURE were performed on the same LD-pruned SNP set. ADMIXTURE (v1.3.0)^121^ was run with K = 2-6 and the optimal K was evaluated using cross-validation errors. Cross-validation (CV) errors were calculated to determine the optimal *K* value. For phylogenetic reconstruction, a pairwise genetic distance matrix was calculated using VCF2Dis (v1.53)^122^ based on the unlinked SNP set, which was subsequently used to construct a Neighbor-Joining (NJ) tree. Crucially, we refined the dataset by excluding individuals with potential signals of recent admixture. Based on the ADMIXTURE results at *K*=3, six individuals exhibiting a dominant ancestry proportion (*Q*) < 0.99 were removed. The remaining 135 individuals (47 *C. grayii*, 43 *C. mystus*, and 45 *C. nasus*), which showed high genetic integrity, constituted the core dataset for evolutionary history reconstruction.

### Genomic diversity and differentiation statistics

Genome-wide nucleotide diversity (π), relative divergence (*F_ST_*), absolute divergence (*D_XY_*), and Tajima’s *D* were estimated using pixy (v2.0.0)^123^ in non-overlapping 1-kb windows using an all-sites VCF masked for accessibility.

### Linkage disequilibrium, gene flow, and local structure

LD decay was quantified using PopLDdecay^124^. Historical relationships and migration edges were explored using TreeMix^125^. Fine-scale heterogeneity in population structure was assessed using winpca^126^ on biallelic SNPs filtered for MAF > 0.05 using a sliding window of 500 kb with a step size of 10 kb.

### Hierarchical demographic inference and speciation modeling

We reconstructed demographic history using a hierarchical strategy combining SMC++ (v1.15.2)^127^ and fastsimcoal2 (v2.8)^128^. To reduce the influence of fine-scale spatial structure, demographic analyses were performed on representative cohorts sampled from single localities: 8 *C. nasus* from the Zhoushan Archipelago (30.73°N, 122.45°E), 12 *C. grayii* from waters adjacent to Qi’ao Island (22.39°N, 113.64°E), and 12 *C. mystus* from waters adjacent to Neilingding Island (22.43°N, 113.83°E) (Supplementary Table S1).

Using these cohorts, we inferred effective population size trajectories and cross-population coalescence rates using SMC++. VCF files were converted using vcf2smc with the mappability mask applied. Uncertainty was assessed via chromosome jackknife resampling^129^ by iteratively excluding one chromosome. Parameters were scaled using a generation time of 1 year and a mutation rate of 2.0 × 10⁻⁹ substitutions per site per generation from Atlantic herring^130^. Finally, global sea-level reconstructions^131^ were used to contextualize the inferred *Ne* variations with paleo-environmental changes.

The multidimensional joint site frequency spectrum was generated using easySFS^132^. from the single-location subsets. We designed and tested four competing scenarios of divergence:

1. Strict Isolation (SI): Species divergence with no subsequent gene flow.
2. Isolation with Migration (IM): Continuous gene flow following divergence.
3. Secondary Contact (SC): An initial isolation period followed by a recent pulse of gene flow.
4. Ancient Gene Flow with Stepwise Isolation (AGF-SI): A complex scenario involving hierarchical divergence, where ancient gene flow occurred between *C. mystus* and the ancestor of *C. grayii*/*C. nasus*, followed by a gradual, stepwise cessation of migration channels.

To enhance model identifiability, the search space for time parameters in fastsimcoal2 was constrained using informative priors: divergence times (*T_div_*) were bounded by MCMCTree phylogenetic estimates, and gene flow cessation times (*T_stop_*) were constrained by the 95% confidence intervals of split times estimated from the 23 SMC++ jackknife replicates. For each model, 100 independent runs (500,000 simulations, 100 EM cycles) were performed. The best-fitting model was selected based on the maximum likelihood and the lowest Akaike Information Criterion (AIC) values. Confidence intervals for parameters were estimated via 100 parametric bootstraps.

### Paleogeographic reconstruction and visualization

To visualize the paleogeographic context, we reconstructed the East Asian margin paleogeography from 30-1 Ma using 6 arc-minute PaleoDEMs from the PALEOMAP Project^133,134^. Map rendering was performed in R using terra^135^ with hypsometric tinting by elevation classes: plains (<200 m), hills (200-500 m), mountains (500-1,000 m), and plateaus (>1,000 m). Paleo-oceanographic features (including the Paleo-Kuroshio Current and Circum-Polar Deep Water) and hypothesized migration routes were overlaid following published reconstructions^136^.

### Detection of adaptive introgression via supervised machine learning

We identified introgressed genomic regions using FILET^137^. Training datasets were generated using coalescent simulations in msmove^138,139^ parameterized using fastsimcoal2 estimates under the best-supported model. We simulated 2 × 10⁴ replicates per class and trained an Extra-Trees classifier to assign genomic windows to three classes: no introgression, introgression from species 1 to species 2, and introgression from species 2 to species 1.

Empirical summary statistics were computed in non-overlapping 1-kb windows across autosomes. Genotypes were phased using SHAPEIT4 (v4.2)^140^. Summary statistics followed Schrider et al.^137^, we incorporated multiple summary statistics to enhance model sensitivity. These included the following within-species statistics (calculated for both source and recipient populations): nucleotide diversity (*pi*), Tajima’s *D*, density of segregating sites (*S*), variance in pairwise distances (hetVar), and the percentile ranking of the minimum pairwise divergence (*dd-*Rank). The following between-species statistics were used: *F_ST_*, absolute genetic divergence (*D_xy_*), minimum pairwise distance (*d_min_*), *G_min_* (defined as *d_min_*/*d_xy_*), and the Nearest-neighbor statistic (*S_nn_*)^141^.

To ensure the robustness of identified introgressed regions, we applied a stringent post-inference filtering strategy. First, empirical windows were classified using the trained classifier, and only those with a posterior probability ≥ 0.99 were retained. Second, adjacent significant windows within 2 kb were merged into single candidate regions. Finally, to distinguish adaptive introgression from neutral gene flow or artifacts, we required candidate regions to overlap with independent signatures of positive selection—specifically, the top 5% outliers of the integrated Haplotype Score (iHS) and number of segregating sites by length (nSL) calculated using selscan (v2.0)^142^.

### Transcriptomic profiling and analysis of regulatory divergence at introgressed loci

To examine regulatory divergence associated with introgressed loci, we analyzed muscle RNA-seq from four wild population groups (n = 4 per group) spanning a hydrological gradient: freshwater lower Yangtze River (*C. nasus*), semi-enclosed Sansha Bay (North *C. mystus*), and the dynamic Pearl River Estuary (South *C. mystus* and *C. grayii*). Strand-specific libraries for *C. mystus* (North/South) and *C. grayii* were sequenced on DNBSEQ-T7 (150-bp paired-end). Raw RNA-seq reads for *C. nasus* were obtained from NCBI SRA (BioProject PRJNA1273475). Sample metadata and mapping statistics are reported in Supplementary Table S3.

All datasets were processed with a uniform pipeline to limit technical heterogeneity. Reads were quality-filtered with fastp (v0.23.0) and aligned to the *C. mystus* reference genome using STAR (v2.7.10a)^143^. Because cross-species alignment can introduce mapping bias, we evaluated alignment performance for each library; all libraries showed robust mapping (Total mapping rate > 94%) and were retained for downstream analyses (Supplementary Table S3). Gene-level read counts were quantified using featureCounts (v2.0.1)^144^. To account for multi-mapping reads caused by gene duplication in teleost genomes, we enabled fractional assignment of multi-mapping reads (-M --fraction).

Raw count matrices were imported into R for downstream analyses. Genes with zero counts across all samples were removed. Differential expression analyses were performed using DESeq2 (v1.50.2)^145^ under a negative binomial generalized linear model. For global visualization, count data were variance-stabilized (VST) to generate principal component analysis (PCA) plots.

Candidate genes were defined a priori as genes overlapping FILET introgression windows (posterior probability ≥ 0.99) and, for adaptive candidates, additionally intersecting the top 5% tails of iHS and nSL. For each introgression direction, expression was compared among recipient, putative donor, and non-introgressed sister lineages following the inferred directionality. Regulatory outcomes were assigned using explicit thresholds based on DESeq2 contrasts (|log_2_FC| > 1; *P_adj_* < 0.05): conserved (recipient not significantly different from sister), convergent (recipient differs from sister but not from donor), or transgressive (recipient differs from both donor and sister), with directionality annotated as transgressive up- or downregulation by the sign of log_2_FC.

## Data availability

The raw whole-genome re-sequencing data and transcriptomic expression data generated in this study have been deposited in the CNGB Sequence Archive (CNSA) of China National GeneBank DataBase (CNGBdb) under project accession number CNP0009055. The *C. mystus* reference genome assembly generated in this study has been deposited at GenBank under accession number GCA_050626335.1. Genome annotations (GFF3), coding sequences (CDS), and protein sequences are available on Figshare at https://doi.org/10.6084/m9.figshare.31342996. The raw sequencing data used for genome assembly (including PacBio HiFi, Hi-C, and RNA-seq reads) are available in the NCBI Sequence Read Archive (SRA) under BioProject accession number PRJNA1161598.

## Acknowledgements

This work was supported by the Innovation Group Project of Southern Marine Science and Engineering Guangdong Laboratory (Zhuhai) (Grant No. 311021006), the R&D Project for Jinwan Yellowfin Seabream Breeding System Construction (Award No. K20-42000-018), and the National Natural Science Foundation of China (Grant No. 31902427).

## Author Contributions

- Conceptualization: Z.F., J.L., B.B.
- Data curation: Z.F., H.W.
- Formal analysis: Z.F., H.W., Y.F., Q.C.
- Funding acquisition: J.L., B.B.
- Investigation: Z.F., Y.H., Z.S., Z.G., W.Z., X.L.
- Methodology: Z.F., H.W., J.L.
- Project administration: J.L.
- Resources: J.L., B.B.
- Software: Z.F., H.W., Y.H.
- Supervision: J.L., B.B.
- Validation: Z.F., H.W., W.Z., X.L., S.L.
- Visualization: Z.F.
- Writing – original draft: Z.F.
- Writing – review & editing: Z.F., H.W., J.L., B.B.

