## Supplementary Information for "Geoclimatic oscillations and ancient reciprocal adaptive introgression shape the evolutionary trajectories of threatened *Coilia*"

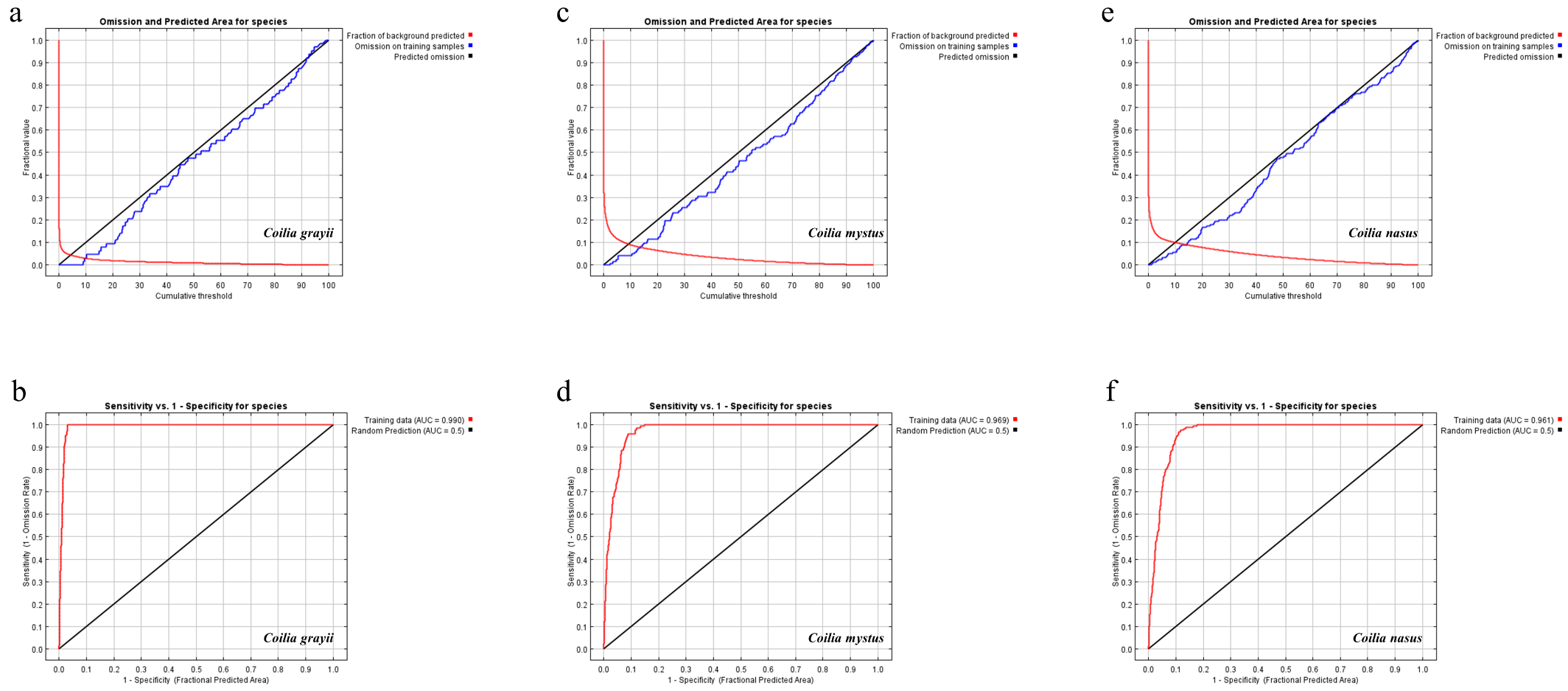

Figure S 1 Performance evaluation of MaxEnt models for *Coilia* species. Receiver operating characteristic (ROC) curves (left panels) and omission/commission error plots (right panels) are shown for **a, b,** *C. grayii* (AUC = 0.990); **c, d,** *C. mystus* (AUC = 0.969); and **e, f,** *C. nasus* (AUC = 0.961). In the ROC plots (**b, d, f**), the red line represents model performance on training data, while the diagonal black line indicates random prediction (AUC = 0.5). High AUC values across all lineages indicate strong model discriminatory power. In the omission plots (**a, c, e**), the close alignment between the training omission rate (blue line) and the predicted omission rate (black line) suggests that the models are well-calibrated and robust against overfitting.

Table S 1 Detailed sampling information and genomic alignment statistics for the *Coilia* individuals analyzed in this study. The table lists the species identity, specific sampling location, habitat type, geographic coordinates, data source, and the percentage of reads successfully mapped to the *C. mystus* reference genome (Mapping Rate) for each individual.

| Sample ID | Species | Sampling Location | Habitat Type | Latitude (N) | Longitude (E) | Data Source | Mapping Rate |
| --- | --- | --- | --- | --- | --- | --- | --- |
| PHC1 | *C. nasus* | Poyang Lake, Jiangxi | Freshwater | 29.4900 | 116.0200 | This study | 98.90% |
| PHC2 | *C. nasus* | Poyang Lake, Jiangxi | Freshwater | 29.4900 | 116.0200 | This study | 98.71% |
| PHC3 | *C. nasus* | Poyang Lake, Jiangxi | Freshwater | 29.4900 | 116.0200 | This study | 98.26% |
| PHC4 | *C. nasus* | Poyang Lake, Jiangxi | Freshwater | 29.4900 | 116.0200 | This study | 98.71% |
| PHC5 | *C. nasus* | Poyang Lake, Jiangxi | Freshwater | 29.4900 | 116.0200 | This study | 98.69% |
| PHC6 | *C. nasus* | Poyang Lake, Jiangxi | Freshwater | 29.4900 | 116.0200 | This study | 98.89% |
| PHC7 | *C. nasus* | Poyang Lake, Jiangxi | Freshwater | 29.4900 | 116.0200 | This study | 98.78% |
| PHC8 | *C. nasus* | Poyang Lake, Jiangxi | Freshwater | 29.4900 | 116.0200 | This study | 98.52% |
| SC1 | *C. nasus* | Zhoushan, Zhejiang | Marine | 30.7300 | 122.4500 | This study | 98.51% |
| SC2 | *C. nasus* | Zhoushan, Zhejiang | Marine | 30.7300 | 122.4500 | This study | 98.38% |
| SC3 | *C. nasus* | Zhoushan, Zhejiang | Marine | 30.7300 | 122.4500 | This study | 98.30% |
| SC4 | *C. nasus* | Zhoushan, Zhejiang | Marine | 30.7300 | 122.4500 | This study | 98.21% |
| SC5 | *C. nasus* | Zhoushan, Zhejiang | Marine | 30.7300 | 122.4500 | This study | 98.49% |
| SC6 | *C. nasus* | Zhoushan, Zhejiang | Marine | 30.7300 | 122.4500 | This study | 98.47% |
| SC7 | *C. nasus* | Zhoushan, Zhejiang | Marine | 30.7300 | 122.4500 | This study | 98.59% |
| SC8 | *C. nasus* | Zhoushan, Zhejiang | Marine | 30.7300 | 122.4500 | This study | 98.48% |
| AP1 | *C. nasus* | Yangzhou (Yangtze River) | Freshwater | 32.2274 | 119.3643 | PRJNA7440151 | 98.88% |
| AP2 | *C. nasus* | Yangzhou (Yangtze River) | Freshwater | 32.2274 | 119.3643 | PRJNA7440151 | 98.92% |
| AP3 | *C. nasus* | Yangzhou (Yangtze River) | Freshwater | 32.2274 | 119.3643 | PRJNA7440151 | 98.83% |
| AP4 | *C. nasus* | Yangzhou (Yangtze River) | Freshwater | 32.2274 | 119.3643 | PRJNA7440151 | 98.73% |
| AP5 | *C. nasus* | Yangzhou (Yangtze River) | Freshwater | 32.2274 | 119.3643 | PRJNA7440151 | 98.86% |
| AP6 | *C. nasus* | Yangzhou (Yangtze River) | Freshwater | 32.2274 | 119.3643 | PRJNA7440151 | 98.84% |
| AP7 | *C. nasus* | Yangzhou (Yangtze River) | Freshwater | 32.2274 | 119.3643 | PRJNA7440151 | 98.72% |
| AP8 | *C. nasus* | Yangzhou (Yangtze River) | Freshwater | 32.2274 | 119.3643 | PRJNA7440151 | 98.52% |
| AP9 | *C. nasus* | Yangzhou (Yangtze River) | Freshwater | 32.2274 | 119.3643 | PRJNA7440151 | 98.89% |
| AP10 | *C. nasus* | Yangzhou (Yangtze River) | Freshwater | 32.2274 | 119.3643 | PRJNA7440151 | 98.96% |
| LP1 | *C. nasus* | Taihu Lake | Freshwater | 31.3271 | 120.0245 | PRJNA7440151 | 98.88% |
| LP2 | *C. nasus* | Taihu Lake | Freshwater | 31.3271 | 120.0245 | PRJNA7440151 | 98.86% |
| LP3 | *C. nasus* | Taihu Lake | Freshwater | 31.3271 | 120.0245 | PRJNA7440151 | 98.87% |
| LP4 | *C. nasus* | Taihu Lake | Freshwater | 31.3271 | 120.0245 | PRJNA7440151 | 98.95% |
| LP5 | *C. nasus* | Taihu Lake | Freshwater | 31.3271 | 120.0245 | PRJNA7440151 | 98.98% |
| LP6 | *C. nasus* | Taihu Lake | Freshwater | 31.3271 | 120.0245 | PRJNA7440151 | 98.91% |
| LP7 | *C. nasus* | Taihu Lake | Freshwater | 31.3271 | 120.0245 | PRJNA7440151 | 98.95% |
| LP8 | *C. nasus* | Taihu Lake | Freshwater | 31.3271 | 120.0245 | PRJNA7440151 | 98.93% |
| LP9 | *C. nasus* | Taihu Lake | Freshwater | 31.3271 | 120.0245 | PRJNA7440151 | 98.91% |
| LP10 | *C. nasus* | Taihu Lake | Freshwater | 31.3271 | 120.0245 | PRJNA7440151 | 98.74% |
| SP1 | *C. nasus* | East China Sea | Marine | 31.8646 | 122.5728 | PRJNA7440151 | 98.94% |
| SP2 | *C. nasus* | East China Sea | Marine | 31.8646 | 122.5728 | PRJNA7440151 | 99.04% |
| SP3 | *C. nasus* | East China Sea | Marine | 31.8646 | 122.5728 | PRJNA7440151 | 99.00% |
| SP4 | *C. nasus* | East China Sea | Marine | 31.8646 | 122.5728 | PRJNA7440151 | 99.14% |
| SP5 | *C. nasus* | East China Sea | Marine | 31.8646 | 122.5728 | PRJNA7440151 | 98.25% |
| SP6 | *C. nasus* | East China Sea | Marine | 31.8646 | 122.5728 | PRJNA7440151 | 98.99% |
| SP7 | *C. nasus* | East China Sea | Marine | 31.8646 | 122.5728 | PRJNA7440151 | 98.92% |
| SP8 | *C. nasus* | East China Sea | Marine | 31.8646 | 122.5728 | PRJNA7440151 | 98.90% |
| SP9 | *C. nasus* | East China Sea | Marine | 31.8646 | 122.5728 | PRJNA7440151 | 98.95% |
| SP10 | *C. nasus* | East China Sea | Marine | 31.8646 | 122.5728 | PRJNA7440151 | 98.99% |
| NDJ1 | *C. mystus* | Sansha Bay (Ningde) | Marine | 26.6942 | 119.7161 | This study | 99.32% |
| NDJ2 | *C. mystus* | Sansha Bay (Ningde) | Marine | 26.6942 | 119.7161 | This study | 99.03% |
| NDJ3 | *C. mystus* | Sansha Bay (Ningde) | Marine | 26.6942 | 119.7161 | This study | 99.13% |
| NDJ04 | *C. mystus* | Sansha Bay (Ningde) | Marine | 26.6942 | 119.7161 | This study | 99.30% |
| NDJ5 | *C. mystus* | Sansha Bay (Ningde) | Marine | 26.6942 | 119.7161 | This study | 99.31% |
| NDJ6 | *C. mystus* | Sansha Bay (Ningde) | Marine | 26.6942 | 119.7161 | This study | 99.26% |
| NDJ7 | *C. mystus* | Sansha Bay (Ningde) | Marine | 26.6942 | 119.7161 | This study | 99.51% |
| NDJ8 | *C. mystus* | Sansha Bay (Ningde) | Marine | 26.6942 | 119.7161 | This study | 99.26% |
| NDJ9 | *C. mystus* | Sansha Bay (Ningde) | Marine | 26.6942 | 119.7161 | This study | 99.13% |
| NDJ10 | *C. mystus* | Sansha Bay (Ningde) | Marine | 26.6942 | 119.7161 | This study | 98.34% |
| NDJ11 | *C. mystus* | Sansha Bay (Ningde) | Marine | 26.6942 | 119.7161 | This study | 99.11% |
| NDJ12 | *C. mystus* | Sansha Bay (Ningde) | Marine | 26.6942 | 119.7161 | This study | 99.19% |
| NDJ13 | *C. mystus* | Sansha Bay (Ningde) | Marine | 26.6942 | 119.7161 | This study | 99.13% |
| NDJ14 | *C. mystus* | Sansha Bay (Ningde) | Marine | 26.6942 | 119.7161 | This study | 98.87% |
| NDJ15 | *C. mystus* | Sansha Bay (Ningde) | Marine | 26.6942 | 119.7161 | This study | 98.38% |
| F2J02 | *C. mystus* | Pearl River Estuary | Estuarine | 23.0389 | 113.5444 | This study | 99.52% |
| F2J03 | *C. mystus* | Pearl River Estuary | Estuarine | 23.0389 | 113.5444 | This study | 99.47% |
| F2J04 | *C. mystus* | Pearl River Estuary | Estuarine | 23.0389 | 113.5444 | This study | 99.51% |
| F2J05 | *C. mystus* | Pearl River Estuary | Estuarine | 23.0389 | 113.5444 | This study | 99.50% |
| F2J06 | *C. mystus* | Pearl River Estuary | Estuarine | 23.0389 | 113.5444 | This study | 99.48% |
| F1J01 | *C. mystus* | Pearl River Estuary | Estuarine | 22.7992 | 113.6111 | This study | 99.41% |
| F1J02 | *C. mystus* | Pearl River Estuary | Estuarine | 22.7992 | 113.6111 | This study | 99.44% |
| F1J03 | *C. mystus* | Pearl River Estuary | Estuarine | 22.7992 | 113.6111 | This study | 99.48% |
| F1J04 | *C. mystus* | Pearl River Estuary | Estuarine | 22.7992 | 113.6111 | This study | 99.35% |
| F1J05 | *C. mystus* | Pearl River Estuary | Estuarine | 22.7992 | 113.6111 | This study | 99.52% |
| F1J06 | *C. mystus* | Pearl River Estuary | Estuarine | 22.7992 | 113.6111 | This study | 99.10% |
| F1J07 | *C. mystus* | Pearl River Estuary | Estuarine | 22.7992 | 113.6111 | This study | 99.41% |
| F1J08 | *C. mystus* | Pearl River Estuary | Estuarine | 22.7992 | 113.6111 | This study | 99.51% |
| F1J09 | *C. mystus* | Pearl River Estuary | Estuarine | 22.7992 | 113.6111 | This study | 98.81% |
| F1J10 | *C. mystus* | Pearl River Estuary | Estuarine | 22.7992 | 113.6111 | This study | 99.47% |
| F1J11 | *C. mystus* | Pearl River Estuary | Estuarine | 22.7992 | 113.6111 | This study | 99.35% |
| F1J12 | *C. mystus* | Pearl River Estuary | Estuarine | 22.7992 | 113.6111 | This study | 99.48% |
| F3J01 | *C. mystus* | Neilingding Island | Estuarine | 22.4308 | 113.8261 | This study | 99.49% |
| F3J02 | *C. mystus* | Neilingding Island | Estuarine | 22.4308 | 113.8261 | This study | 99.47% |
| F3J03 | *C. mystus* | Neilingding Island | Estuarine | 22.4308 | 113.8261 | This study | 99.46% |
| F3J04 | *C. mystus* | Neilingding Island | Estuarine | 22.4308 | 113.8261 | This study | 99.49% |
| F3J05 | *C. mystus* | Neilingding Island | Estuarine | 22.4308 | 113.8261 | This study | 99.50% |
| F3J06 | *C. mystus* | Neilingding Island | Estuarine | 22.4308 | 113.8261 | This study | 99.44% |
| F3J07 | *C. mystus* | Neilingding Island | Estuarine | 22.3944 | 113.7889 | This study | 99.48% |
| F3J08 | *C. mystus* | Neilingding Island | Estuarine | 22.3944 | 113.7889 | This study | 99.51% |
| F3J09 | *C. mystus* | Neilingding Island | Estuarine | 22.3944 | 113.7889 | This study | 99.48% |
| F3J10 | *C. mystus* | Neilingding Island | Estuarine | 22.3944 | 113.7889 | This study | 99.53% |
| F3J11 | *C. mystus* | Neilingding Island | Estuarine | 22.3944 | 113.7889 | This study | 99.53% |
| F3J12 | *C. mystus* | Neilingding Island | Estuarine | 22.3944 | 113.7889 | This study | 99.50% |
| QYJ1 | *C. grayii* | Beijiang River (Yingde) | Freshwater | 23.9229 | 113.2833 | This study | 98.83% |
| QYJ2 | *C. grayii* | Beijiang River (Yingde) | Freshwater | 23.9229 | 113.2833 | This study | 98.81% |
| QYJ3 | *C. grayii* | Beijiang River (Yingde) | Freshwater | 23.9368 | 113.2677 | This study | 98.86% |
| QYJ4 | *C. grayii* | Beijiang River (Yingde) | Freshwater | 23.9368 | 113.2677 | This study | 98.88% |
| QYJ5 | *C. grayii* | Beijiang River (Yingde) | Freshwater | 23.9368 | 113.2677 | This study | 98.94% |
| QYJ6 | *C. grayii* | Beijiang River (Yingde) | Freshwater | 23.9123 | 113.2689 | This study | 98.96% |
| QYJ7 | *C. grayii* | Beijiang River (Yingde) | Freshwater | 23.9123 | 113.2689 | This study | 98.89% |
| QYJ8 | *C. grayii* | Beijiang River (Yingde) | Freshwater | 23.9336 | 113.2476 | This study | 98.80% |
| QYJ9 | *C. grayii* | Beijiang River (Yingde) | Freshwater | 23.9336 | 113.2476 | This study | 98.96% |
| QYJ10 | *C. grayii* | Beijiang River (Yingde) | Freshwater | 23.9336 | 113.2476 | This study | 98.90% |
| QYJ11 | *C. grayii* | Beijiang River (Yingde) | Freshwater | 23.9449 | 113.2782 | This study | 98.90% |
| QYJ12 | *C. grayii* | Beijiang River (Yingde) | Freshwater | 23.9449 | 113.2782 | This study | 98.90% |
| QYJ13 | *C. grayii* | Beijiang River (Yingde) | Freshwater | 23.8812 | 113.2602 | This study | 98.86% |
| QYJ14 | *C. grayii* | Beijiang River (Yingde) | Freshwater | 23.8812 | 113.2602 | This study | 98.87% |
| QYJ15 | *C. grayii* | Beijiang River (Yingde) | Freshwater | 23.8812 | 113.2602 | This study | 98.87% |
| Q2J01 | *C. grayii* | Pearl River Estuary | Estuarine | 23.0840 | 113.5542 | This study | 98.92% |
| Q2J02 | *C. grayii* | Pearl River Estuary | Estuarine | 23.0840 | 113.5542 | This study | 98.94% |
| Q2J03 | *C. grayii* | Pearl River Estuary | Estuarine | 23.0840 | 113.5542 | This study | 98.97% |
| Q2J04 | *C. grayii* | Pearl River Estuary | Estuarine | 23.0840 | 113.5542 | This study | 99.06% |
| Q2J05 | *C. grayii* | Pearl River Estuary | Estuarine | 23.0840 | 113.5542 | This study | 98.89% |
| Q2J06 | *C. grayii* | Pearl River Estuary | Estuarine | 23.0840 | 113.5542 | This study | 98.94% |
| Q2J07 | *C. grayii* | Pearl River Estuary | Estuarine | 23.0840 | 113.5542 | This study | 98.98% |
| Q2J08 | *C. grayii* | Pearl River Estuary | Estuarine | 23.0474 | 113.5231 | This study | 98.87% |
| Q2J09 | *C. grayii* | Pearl River Estuary | Estuarine | 23.0474 | 113.5231 | This study | 98.87% |
| Q2J10 | *C. grayii* | Pearl River Estuary | Estuarine | 23.0474 | 113.5231 | This study | 98.88% |
| Q2J11 | *C. grayii* | Pearl River Estuary | Estuarine | 23.0394 | 113.5124 | This study | 98.92% |
| Q2J12 | *C. grayii* | Pearl River Estuary | Estuarine | 23.0394 | 113.5124 | This study | 98.91% |
| Q1J01 | *C. grayii* | Pearl River Estuary | Estuarine | 23.0394 | 113.5124 | This study | 98.95% |
| Q1J02 | *C. grayii* | Pearl River Estuary | Estuarine | 22.8126 | 113.6162 | This study | 98.84% |
| Q1J03 | *C. grayii* | Pearl River Estuary | Estuarine | 22.8126 | 113.6162 | This study | 98.88% |
| Q1J04 | *C. grayii* | Pearl River Estuary | Estuarine | 22.8126 | 113.6162 | This study | 98.90% |
| Q1J05 | *C. grayii* | Pearl River Estuary | Estuarine | 22.8074 | 113.6122 | This study | 98.94% |
| Q1J06 | *C. grayii* | Pearl River Estuary | Estuarine | 22.8074 | 113.6122 | This study | 98.89% |
| Q1J07 | *C. grayii* | Pearl River Estuary | Estuarine | 22.8074 | 113.6122 | This study | 98.91% |
| Q1J08 | *C. grayii* | Pearl River Estuary | Estuarine | 22.7865 | 113.6255 | This study | 98.93% |
| Q1J09 | *C. grayii* | Pearl River Estuary | Estuarine | 22.7865 | 113.6255 | This study | 98.89% |
| Q1J10 | *C. grayii* | Pearl River Estuary | Estuarine | 22.7865 | 113.6255 | This study | 98.92% |
| Q1J11 | *C. grayii* | Pearl River Estuary | Estuarine | 22.4669 | 113.6585 | This study | 98.92% |
| Q1J12 | *C. grayii* | Pearl River Estuary | Estuarine | 22.4669 | 113.6585 | This study | 98.94% |
| Q4J01 | *C. grayii* | Qi’ao Island | Estuarine | 22.4669 | 113.6585 | This study | 98.86% |
| Q4J02 | *C. grayii* | Qi’ao Island | Estuarine | 22.4669 | 113.6585 | This study | 98.90% |
| Q4J03 | *C. grayii* | Qi’ao Island | Estuarine | 22.4132 | 113.6773 | This study | 98.90% |
| Q4J04 | *C. grayii* | Qi’ao Island | Estuarine | 22.4132 | 113.6773 | This study | 98.88% |
| Q4J05 | *C. grayii* | Qi’ao Island | Estuarine | 22.4132 | 113.6773 | This study | 98.87% |
| Q4J06 | *C. grayii* | Qi’ao Island | Estuarine | 22.4132 | 113.6773 | This study | 98.86% |
| Q4J07 | *C. grayii* | Qi’ao Island | Estuarine | 22.4132 | 113.6773 | This study | 98.88% |
| Q4J08 | *C. grayii* | Qi’ao Island | Estuarine | 22.4132 | 113.6773 | This study | 98.87% |
| Q4J09 | *C. grayii* | Qi’ao Island | Estuarine | 22.3911 | 113.6449 | This study | 98.84% |
| Q4J10 | *C. grayii* | Qi’ao Island | Estuarine | 22.3911 | 113.6449 | This study | 98.89% |
| Q4J11 | *C. grayii* | Qi’ao Island | Estuarine | 22.3911 | 113.6449 | This study | 98.96% |
| Q4J12 | *C. grayii* | Qi’ao Island | Estuarine | 22.3911 | 113.6449 | This study | 98.96% |

Note: To mitigate potential confounding effects arising from spatial population substructure, SC1-SC8, F3J01-F3J12, Q4J01-Q4J12 were used for demographic inference.

Table S 2 Statistics of sequencing depth supporting the ZZ/ZO sex determination system. sequencing depth was calculated using Mosdepth and normalized by the average depth of all autosomes. Male samples (n=2; F1J09, F3J10) show a normalized depth >1.0 on ChrZ, while female samples (n=28) show reduced coverage (~0.68), resulting in an M/F ratio of 1.87, consistent with a ZZ/ZO sex-determination system.

| Chromosome | Original Scaffold | Size(bp) | Norm. Depth (Male, n=2) | Norm. Depth (Female, n=28) | M/F Ratio | Inference |
| --- | --- | --- | --- | --- | --- | --- |
| Chr01 | R5 | 41143600 | 1.03 | 1 | 1.03 | Autosome |
| Chr02 | R2 | 39957351 | 0.96 | 0.99 | 0.97 | Autosome |
| Chr03 | R3 | 38667571 | 1.05 | 1.04 | 1.01 | Autosome |
| Chr04 | R4 | 38338110 | 1.07 | 1.07 | 1 | Autosome |
| Chr05 | R10 | 37359565 | 1.01 | 1.01 | 1 | Autosome |
| Chr06 | R6 | 37128509 | 1.09 | 1.1 | 0.99 | Autosome |
| Chr07 | R7 | 37046970 | 1.08 | 1.08 | 1 | Autosome |
| Chr08 | R8 | 36023233 | 1.05 | 1.06 | 0.99 | Autosome |
| Chr09 | R9 | 35872567 | 0.95 | 0.94 | 1.01 | Autosome |
| Chr10 | R13 | 35377138 | 1.07 | 1.07 | 1 | Autosome |
| Chr11 | R11 | 34841370 | 1.02 | 1.03 | 0.99 | Autosome |
| Chr12 | R12 | 34459153 | 1.03 | 1.03 | 1 | Autosome |
| Chr13 | R18 | 34318340 | 1.03 | 1.03 | 1 | Autosome |
| Chr14 | R16 | 33971454 | 1.03 | 1.05 | 0.98 | Autosome |
| Chr15 | R14 | 33946335 | 1.03 | 1.04 | 0.99 | Autosome |
| Chr16 | R19 | 33874442 | 1.03 | 1.04 | 0.99 | Autosome |
| Chr17 | R15 | 33602267 | 1.02 | 1.02 | 1 | Autosome |
| Chr18 | R21 | 33092659 | 1.06 | 1.06 | 1 | Autosome |
| Chr19 | R20 | 32389169 | 1.06 | 1.07 | 0.99 | Autosome |
| Chr20 | R17 | 31418414 | 0.94 | 0.94 | 1 | Autosome |
| Chr21 | R22 | 30576199 | 0.98 | 0.99 | 0.99 | Autosome |
| Chr22 | R24 | 29603253 | 0.98 | 0.99 | 0.99 | Autosome |
| Chr23 | R23 | 28342729 | 0.99 | 0.98 | 1.01 | Autosome |
| **ChrZ** | **R1** | **45142098** | **1.27** | **0.68** | **1.87** | **Sex Chr (Z)** |

Table S 3 Summary of transcriptomic samples, sequencing quality, and mapping statistics. (Note: All samples were derived from dorsal muscle tissues. Sequencing quality was assessed by the Q30 percentage of clean reads. Total mapping rate includes uniquely mapped reads and multi-mapping reads retained by fractional counting.)

| Sample ID | Species | Group | Sampling Location | Seq Platform | Clean Reads (M) | Q30 (%) | Total Mapping Rate (%)† | Data Source |
| --- | --- | --- | --- | --- | --- | --- | --- | --- |
| F3J1 | *C. mystus* | South_CMY | Pearl River Estuary | DNBSEQ-T7 | 52.42 | 94.83 | 98.2 | This study |
| F3J2 | *C. mystus* | South_CMY | Pearl River Estuary | DNBSEQ-T7 | 50.24 | 94.47 | 97.6 | This study |
| F3J3 | *C. mystus* | South_CMY | Pearl River Estuary | DNBSEQ-T7 | 44.71 | 94.84 | 98.4 | This study |
| F3J4 | *C. mystus* | South_CMY | Pearl River Estuary | DNBSEQ-T7 | 50.37 | 94.74 | 98.2 | This study |
| NDJ1 | *C. mystus* | North_CMY | Sansha Bay (Ningde) | DNBSEQ-T7 | 45.26 | 94.32 | 98.5 | This study |
| NDJ2 | *C. mystus* | North_CMY | Sansha Bay (Ningde) | DNBSEQ-T7 | 38.11 | 93.82 | 88.2 | This study |
| NDJ3 | *C. mystus* | North_CMY | Sansha Bay (Ningde) | DNBSEQ-T7 | 47.34 | 95.80 | 92.8 | This study |
| NDJ4 | *C. mystus* | North_CMY | Sansha Bay (Ningde) | DNBSEQ-T7 | 42.46 | 94.36 | 96.2 | This study |
| Q4J01 | *C. grayii* | CGR | Pearl River Estuary | DNBSEQ-T7 | 50.51 | 90.86 | 93.1 | This study |
| Q4J02 | *C. grayii* | CGR | Pearl River Estuary | DNBSEQ-T7 | 51.79 | 91.52 | 94.7 | This study |
| Q4J03 | *C. grayii* | CGR | Pearl River Estuary | DNBSEQ-T7 | 51.39 | 90.31 | 91.3 | This study |
| Q4J04 | *C. grayii* | CGR | Pearl River Estuary | DNBSEQ-T7 | 50.31 | 90.50 | 94.2 | This study |
| DJ13 | *C. nasus* | CNA | Yangtze River | Illumina NovaSeq | 36.22 | 95.57 | 96.5 | SRR33880213 |
| DJ14 | *C. nasus* | CNA | Yangtze River | Illumina NovaSeq | 46.09 | 95.92 | 96.1 | SRR33880214 |
| DJ15 | *C. nasus* | CNA | Yangtze River | Illumina NovaSeq | 35.63 | 95.73 | 92.7 | SRR33880215 |
| DJ16 | *C. nasus* | CNA | Yangtze River | Illumina NovaSeq | 43.06 | 95.92 | 94.5 | SRR33880216 |

**Footnotes:** † Group assignments correspond to the labels used in the PCA and differential expression analyses (Fig. 6). † Represents the sum of uniquely mapped reads and multi-mapping reads, quantified using the fractional counting strategy.

Table S 4 Niche overlap estimates and statistical significance for three *Coilia* species across five environmental dimensions.

| Variable | Comparison | Schoener’s *D* | *p_Eq_* | *p_Sim_* |
| --- | --- | --- | --- | --- |
| Temperature | *C. grayii* vs *C. nasus* | 0.116 | 1 | 0.5834 |
|  | *C. grayii* vs *C. mystus* | 0.369 | 1 | 0.3437 |
|  | *C. nasus* vs *C. mystus* | 0.698 | 1 | 0.3886 |
| Depth | *C. grayii* vs *C. nasus* | 0.606 | 1 | **0.013*** |
|  | *C. grayii* vs *C. mystus* | 0.531 | 1 | **0.022*** |
|  | *C. nasus* vs *C. mystus* | 0.846 | 0.8162 | **0.003*** |
| Salinity | *C. grayii* vs *C. nasus* | 0.933 | 0.1738 | **0.001*** |
|  | *C. grayii* vs *C. mystus* | 0.569 | 0.997 | **0.003*** |
|  | *C. nasus* vs *C. mystus* | 0.604 | 1 | **0.043*** |
| Dist_to_coast | *C. grayii* vs *C. nasus* | 0.275 | 0.8951 | 0.0949 |
|  | *C. grayii* vs *C. mystus* | 0.12 | 1 | 0.3437 |
|  | *C. nasus* vs *C. mystus* | 0.097 | 1 | 0.1429 |
| Runoff | *C. grayii* vs *C. nasus* | 0.552 | 1 | 0.7303 |
|  | *C. grayii* vs *C. mystus* | 0.452 | 1 | 0.7632 |
|  | *C. nasus* vs *C. mystus* | 0.889 | 0.3457 | 0.3966 |

Note: Schoener’s *D* was used to quantify niche overlap, ranging from 0 (no overlap) to 1 (identical niches). Statistical significance was assessed using 1,000 permutations within a PCA-env framework, where a *p_Eq_* value approaching 1.000 indicates significant niche divergence (*D_obs_* < *D_null_*), and values within the central 95% interval (0.05 < *p_Eq_* < 0.95) support niche equivalency. For the similarity test, *p_Sim_* < 0.05 (indicated by bold text and asterisks) denotes niches that are significantly more similar than expected by chance. Dist_to_coast refers to the signed distance to the coastline (positive: inland/riverine; negative: offshore/marine), and Runoff represents the surface freshwater discharge at occurrence localities.

Table S 5 Model selection statistics for the four demographic scenarios tested using fastsimcoal2

| Model Scenario | Abbreviation | No. Parameters (k) | ΔLikelihood | AIC | ΔAIC | Support |
| --- | --- | --- | --- | --- | --- | --- |
| Ancient Gene Flow with Stepwise Isolation | AGF-SI | 18 | 518,477 | 129,700,595 | 0 | Best Fit |
| Secondary Contact | SC | 14 | 539,537 | 129,797,811 | 97,216 | - |
| Strict Isolation | SI | 7 | 810,254 | 131,044,495 | 1,343,900 | - |
| Isolation with Migration | IM | 13 | 3,416,588 | 143,047,121 | 13,346,526 | - |

Table S 6 Functional classification of adaptive introgression candidates across *Coilia* lineages. The table summarizes the number of candidate genes associated with specific biological categories for each introgression direction (Donor to Recipient). Gene functions were annotated based on Gene Ontology (GO) and literature mining. "Total" indicates the number of high-confidence adaptive introgressed genes identified by FILET and selection scans (nSL/iHS) in each directional pair.

| Category | CGR to CMY | CMY to CGR | CMY to CNA | CGR to CNA | CNA to CMY | CNA to CGR |
| --- | --- | --- | --- | --- | --- | --- |
| Immunity & Defense | 4 | 24 | 14 | 1 | 0 | 1 |
| Osmoregulation & Ion Transport | 4 | 0 | 6 | 4 | 0 | 0 |
| Muscle, Cytoskeleton & Adhesion | 2 | 9 | 16 | 2 | 0 | 1 |
| Metabolism & Mitochondria | 12 | 5 | 6 | 1 | 5 | 1 |
| Nervous & Sensory System | 4 | 1 | 3 | 3 | 1 | 0 |
| Reproduction | 0 | 3 | 0 | 0 | 0 | 0 |
| Cellular Processes & Regulation | 14 | 16 | 37 | 1 | 3 | 0 |
| Others/Unknown | 10 | 11 | 19 | 1 | 0 | 0 |
| Total | 50 | 68 | 101 | 14 | 9 | 3 |

Table S 7 Detailed annotation and functional description of adaptive introgression candidates identified by combinatorial genomic scans. Candidates were identified using the FILET classifier (posterior probability ≥ 0.99) and validated by selective sweep signals (top 5% of nSL or iHS). Columns provide the direction of introgression, gene identifiers, broad functional categories (corresponding to Supplementary Table 6), and putative functions derived from orthologous annotation in zebrafish (*Danio rerio*) or human databases.

| Direction | Gene ID | Gene Symbol | Broad Category | Function / Description |
| --- | --- | --- | --- | --- |
| CGR2CMY | *Acrfas012729* | *C3* | Immunity & Defense | Complement C3 |
| CGR2CMY | *Acrfas002446* | *CCL34* | Immunity & Defense | Chemokine (C-C motif) ligand 34 |
| CGR2CMY | *Acrfas010656* | *OSMR* | Immunity & Defense | Oncostatin M receptor |
| CGR2CMY | *Acrfas004247* | *TNFRSF* | Immunity & Defense | TNF receptor superfamily member |
| CGR2CMY | *Acrfas016712* | *AQP10* | Osmoregulation & Ion Transport | Aquaporin 10 (Water transport) |
| CGR2CMY | *Acrfas004896* | *KCNH6* | Osmoregulation & Ion Transport | Potassium voltage-gated channel |
| CGR2CMY | *Acrfas018464* | *SLC16A9* | Osmoregulation & Ion Transport | Monocarboxylic acid transporter |
| CGR2CMY | *Acrfas023470* | *SLC35A5* | Osmoregulation & Ion Transport | Solute carrier family 35 |
| CGR2CMY | *Acrfas003238* | *ERC1* | Muscle, Cytoskeleton & Adhesion | ELKS/RAB6-interacting/CAST family |
| CGR2CMY | *Acrfas004260* | *SNTB1* | Muscle, Cytoskeleton & Adhesion | Syntrophin, basic 1 |
| CGR2CMY | *Acrfas025421* | *ASNS* | Metabolism & Mitochondria | Asparagine synthetase |
| CGR2CMY | *Acrfas007909* | *ATG* | Metabolism & Mitochondria | Autophagy-related protein |
| CGR2CMY | *Acrfas022631* | *DHODH* | Metabolism & Mitochondria | Dihydroorotate dehydrogenase |
| CGR2CMY | *Acrfas018303* | *EIF3A* | Metabolism & Mitochondria | Translation initiation factor 3 |
| CGR2CMY | *Acrfas014275* | *ERO1L* | Metabolism & Mitochondria | Oxidative folding (ER stress) |
| CGR2CMY | *Acrfas010391* | *Glycosyltransferase* | Metabolism & Mitochondria | Glycosyltransferase family 92 |
| CGR2CMY | *Acrfas005989* | *ILVBL* | Metabolism & Mitochondria | Acetolactate synthase-like |
| CGR2CMY | *Acrfas003910* | *IREB2* | Metabolism & Mitochondria | Iron-responsive element binding protein 2 |
| CGR2CMY | *Acrfas013867* | *PSMB7* | Metabolism & Mitochondria | Proteasome subunit beta 7 |
| CGR2CMY | *Acrfas003745* | *SLC2A13* | Metabolism & Mitochondria | Glucose transporter (GLUT13) |
| CGR2CMY | *Acrfas012629* | *SULT1* | Metabolism & Mitochondria | Sulfotransferase 1 family |
| CGR2CMY | *Acrfas008565* | *Ssu-2* | Metabolism & Mitochondria | Ssu-2 homolog |
| CGR2CMY | *Acrfas007755* | *ARR3A* | Nervous & Sensory System | Arrestin 3a, retinal |
| CGR2CMY | *Acrfas007086* | *AUTS2* | Nervous & Sensory System | Autism susceptibility gene 2 |
| CGR2CMY | *Acrfas007763* | *OPSIN* | Nervous & Sensory System | Teleost multiple tissue opsin |
| CGR2CMY | *Acrfas020004* | *SEA domain* | Nervous & Sensory System | SEA domain containing protein |
| CGR2CMY | *Acrfas007409* | *AKAP10* | Cellular Processes & Regulation | A-kinase anchor protein 10 |
| CGR2CMY | *Acrfas021853* | *DUSP23* | Cellular Processes & Regulation | Dual specificity phosphatase 23 |
| CGR2CMY | *Acrfas012840* | *ILF3* | Cellular Processes & Regulation | Interleukin enhancer binding factor 3 |
| CGR2CMY | *Acrfas015502* | *KDR* | Cellular Processes & Regulation | VEGFR (Kinase insert domain receptor) |
| CGR2CMY | *Acrfas026439* | *MKL1* | Cellular Processes & Regulation | Megakaryoblastic leukemia 1 |
| CGR2CMY | *Acrfas016064* | *ORC1* | Cellular Processes & Regulation | Origin recognition complex |
| CGR2CMY | *Acrfas014290* | *PKC-CalB* | Cellular Processes & Regulation | Protein kinase C conserved region |
| CGR2CMY | *Acrfas022472* | *PPIP5K1* | Cellular Processes & Regulation | Histidine acid phosphatase |
| CGR2CMY | *Acrfas015743* | *PRKACB* | Cellular Processes & Regulation | Protein kinase, cAMP-dependent |
| CGR2CMY | *Acrfas016850* | *PRSS2* | Cellular Processes & Regulation | Peptidase S1 family |
| CGR2CMY | *Acrfas004860* | *PSMC3IP* | Cellular Processes & Regulation | PSMC3 interacting protein |
| CGR2CMY | *Acrfas026409* | *RNF40* | Cellular Processes & Regulation | Ring finger protein 40 |
| CGR2CMY | *Acrfas003475* | *RUVBL2* | Cellular Processes & Regulation | Chromatin remodeling (INO80 complex) |
| CGR2CMY | *Acrfas006473* | *USP16* | Cellular Processes & Regulation | Ubiquitin specific peptidase 16 |
| CGR2CMY | *Acrfas025860* | *AKHR* | Others/Unknown | G-protein coupled receptor |
| CGR2CMY | *Acrfas002148* | *C18orf8* | Others/Unknown | Chromosome 18 open reading frame 8 |
| CGR2CMY | *Acrfas003508* | *C3orf20* | Others/Unknown | Protein C3orf20 homolog |
| CGR2CMY | *Acrfas020520* | *CIDEB* | Others/Unknown | Cell death-inducing DFFA-like effector b |
| CGR2CMY | *Acrfas019024* | *MANSC1* | Others/Unknown | MANSC domain containing 1 |
| CGR2CMY | *Acrfas009165* | *PRICKLE2* | Others/Unknown | Prickle-like 2 |
| CGR2CMY | *Acrfas006474* | *RWDD2B* | Others/Unknown | RWD domain containing 2B |
| CGR2CMY | *Acrfas021856* | *SELL* | Others/Unknown | AhpC/TSA antioxidant enzyme |
| CGR2CMY | *Acrfas021857* | *SELL* | Others/Unknown | AhpC/TSA antioxidant enzyme |
| CGR2CMY | *Acrfas018030* | *WDR27* | Others/Unknown | WD repeat domain 27 |
| CMY2CGR | *Acrfas023090* | *AIG1* | Immunity & Defense | AIG1 family |
| CMY2CGR | *Acrfas020060* | *APOL* | Immunity & Defense | Apolipoprotein L |
| CMY2CGR | *Acrfas011869* | *BCR-like* | Immunity & Defense | B-cell receptor CD22-like |
| CMY2CGR | *Acrfas019206* | *C-type Lectin* | Immunity & Defense | CTL / CRD domain |
| CMY2CGR | *Acrfas011766* | *GBP1* | Immunity & Defense | Guanylate-binding protein 1 |
| CMY2CGR | *Acrfas022987* | *GTPase IMAP* | Immunity & Defense | GTPase IMAP family member |
| CMY2CGR | *Acrfas005296* | *IFI44* | Immunity & Defense | Interferon-induced protein 44-like |
| CMY2CGR | *Acrfas011767* | *IFI44* | Immunity & Defense | Interferon-induced protein 44-like |
| CMY2CGR | *Acrfas011765* | *IFI44* | Immunity & Defense | Interferon-induced protein 44-like |
| CMY2CGR | *Acrfas004272* | *ITGAD* | Immunity & Defense | Integrin alpha |
| CMY2CGR | *Acrfas011553* | *Ig V-set* | Immunity & Defense | Immunoglobulin V-set |
| CMY2CGR | *Acrfas009950* | *MHC I* | Immunity & Defense | MHC class I |
| CMY2CGR | *Acrfas009921* | *MHC II* | Immunity & Defense | HLA-DPB1 (Antigen presentation) |
| CMY2CGR | *Acrfas012008* | *NACHT* | Immunity & Defense | NACHT, LRR and PYD domains |
| CMY2CGR | *Acrfas011785* | *NLRP12* | Immunity & Defense | Inflammasome / NACHT domain |
| CMY2CGR | *Acrfas011784* | *NLRP12* | Immunity & Defense | Inflammasome / NACHT domain |
| CMY2CGR | *Acrfas022406* | *SEMA7A* | Immunity & Defense | Semaphorin 7A |
| CMY2CGR | *Acrfas011928* | *SIGLEC1* | Immunity & Defense | Sialoadhesin (Macrophage receptor) |
| CMY2CGR | *Acrfas011929* | *SIGLEC1* | Immunity & Defense | Sialoadhesin |
| CMY2CGR | *Acrfas011870* | *SIGLEC1* | Immunity & Defense | Sialic acid binding Ig-like lectin |
| CMY2CGR | *Acrfas008331* | *STAB1* | Immunity & Defense | Stabilin 1 |
| CMY2CGR | *Acrfas023040* | *TRIM35* | Immunity & Defense | Tripartite motif containing 35 |
| CMY2CGR | *Acrfas023131* | *TRIM47* | Immunity & Defense | Tripartite motif containing 47 |
| CMY2CGR | *Acrfas023133* | *TRIM47* | Immunity & Defense | Tripartite motif containing 47 |
| CMY2CGR | *Acrfas005276* | *COL11A1* | Muscle, Cytoskeleton & Adhesion | Collagen type XI alpha 1 |
| CMY2CGR | *Acrfas008790* | *COL7A1* | Muscle, Cytoskeleton & Adhesion | Collagen type VII alpha 1 |
| CMY2CGR | *Acrfas024764* | *DNAH12* | Muscle, Cytoskeleton & Adhesion | Dynein heavy chain 12 |
| CMY2CGR | *Acrfas011665* | *DNAH9* | Muscle, Cytoskeleton & Adhesion | Dynein heavy chain 9 |
| CMY2CGR | *Acrfas005653* | *DSCAML1* | Muscle, Cytoskeleton & Adhesion | Down syndrome cell adhesion |
| CMY2CGR | *Acrfas013436* | *MYH-Fast* | Muscle, Cytoskeleton & Adhesion | Myosin heavy chain, fast |
| CMY2CGR | *Acrfas012847* | *MYH11* | Muscle, Cytoskeleton & Adhesion | Myosin heavy chain 11 |
| CMY2CGR | *Acrfas026584* | *MYOF* | Muscle, Cytoskeleton & Adhesion | Myoferlin |
| CMY2CGR | *Acrfas022505* | *NCAM2* | Muscle, Cytoskeleton & Adhesion | Neural cell adhesion molecule 2 |
| CMY2CGR | *Acrfas015858* | *FTSJ3* | Metabolism & Mitochondria | RNA methyltransferase |
| CMY2CGR | *Acrfas009645* | *GCN1L1* | Metabolism & Mitochondria | GCN1 general control of amino-acid |
| CMY2CGR | *Acrfas003291* | *RT* | Metabolism & Mitochondria | Reverse transcriptase |
| CMY2CGR | *Acrfas010824* | *RT* | Metabolism & Mitochondria | Reverse transcriptase |
| CMY2CGR | *Acrfas010191* | *WDFY3* | Metabolism & Mitochondria | Autophagy linked FYVE protein |
| CMY2CGR | *Acrfas013085* | *C16orf45* | Reproduction | Sperm-associated protein |
| CMY2CGR | *Acrfas011927* | *IZUMO1* | Reproduction | Izumo sperm-egg fusion protein |
| CMY2CGR | *Acrfas026376* | *PLEKHS1* | Reproduction | PLEKHS1 (Sperm acrosome) |
| CMY2CGR | *Acrfas014167* | *AP3B1* | Cellular Processes & Regulation | Adaptor-related protein complex 3 |
| CMY2CGR | *Acrfas026645* | *BAHCC1* | Cellular Processes & Regulation | BAH domain containing 1 |
| CMY2CGR | *Acrfas016318* | *CELA1* | Cellular Processes & Regulation | Chymotrypsin-like elastase 1 |
| CMY2CGR | *Acrfas018063* | *CHUK* | Cellular Processes & Regulation | IKK-alpha (NF-kB pathway) |
| CMY2CGR | *Acrfas002625* | *CNKSR2* | Cellular Processes & Regulation | Connector enhancer of kinase suppressor of ras |
| CMY2CGR | *Acrfas023027* | *CNOT11* | Cellular Processes & Regulation | CCR4-NOT transcription complex |
| CMY2CGR | *Acrfas014359* | *ITSN2* | Cellular Processes & Regulation | Intersectin |
| CMY2CGR | *Acrfas019205* | *MAGI2* | Cellular Processes & Regulation | MAGI2 (Junction protein) |
| CMY2CGR | *Acrfas020259* | *NELFA* | Cellular Processes & Regulation | Negative elongation factor A |
| CMY2CGR | *Acrfas024738* | *NUGGC* | Cellular Processes & Regulation | Nuclear GTPase |
| CMY2CGR | *Acrfas024766* | *PLC-X* | Cellular Processes & Regulation | Phosphatidylinositol-specific PLC |
| CMY2CGR | *Acrfas023951* | *RANBP2* | Cellular Processes & Regulation | RAN binding protein 2 |
| CMY2CGR | *Acrfas005415* | *SAP30BP* | Cellular Processes & Regulation | SAP30 binding protein |
| CMY2CGR | *Acrfas018889* | *SND1* | Cellular Processes & Regulation | Staphylococcal nuclease domain |
| CMY2CGR | *Acrfas020059* | *STK* | Cellular Processes & Regulation | Serine threonine-protein kinase |
| CMY2CGR | *Acrfas002955* | *TOP* | Cellular Processes & Regulation | DNA topoisomerase |
| CMY2CGR | *Acrfas013142* | *DNA-bind* | Others/Unknown | DNA binding |
| CMY2CGR | *Acrfas013128* | *LOXL4* | Others/Unknown | Lysyl oxidase-like 4 |
| CMY2CGR | *Acrfas017576* | *PTTG1IP* | Others/Unknown | Pituitary tumor-transforming gene IP |
| CMY2CGR | *Acrfas026750* | *RNF213* | Others/Unknown | Ring finger protein 213b |
| CMY2CGR | *Acrfas012725* | *Transposon* | Others/Unknown | Plant transposon protein |
| CMY2CGR | *Acrfas005696* | *Uncharacterized* | Others/Unknown | - |
| CMY2CGR | *Acrfas006580* | *Uncharacterized* | Others/Unknown | - |
| CMY2CGR | *Acrfas023319* | *Uncharacterized* | Others/Unknown | - |
| CMY2CGR | *Acrfas018065* | *Uncharacterized* | Others/Unknown | - |
| CMY2CGR | *Acrfas023026* | *Up-regulator* | Others/Unknown | Up-regulator of cell proliferation |
| CMY2CGR | *Acrfas006751* | *Up-regulator* | Others/Unknown | Up-regulator of cell proliferation |
| CMY2CNA | *Acrfas010486* | *CCL21* | Immunity & Defense | Chemokine (C-C motif) ligand 21 |
| CMY2CNA | *Acrfas013937* | *HPSE* | Immunity & Defense | Heparanase |
| CMY2CNA | *Acrfas004272* | *ITGAD* | Immunity & Defense | Integrin alpha |
| CMY2CNA | *Acrfas023120* | *ITIH4* | Immunity & Defense | Inter-alpha-trypsin inhibitor |
| CMY2CNA | *Acrfas011553* | *Ig V-set* | Immunity & Defense | Immunoglobulin V-set |
| CMY2CNA | *Acrfas021725* | *Ig V-set* | Immunity & Defense | Immunoglobulin |
| CMY2CNA | *Acrfas025278* | *MHC I* | Immunity & Defense | MHC class I |
| CMY2CNA | *Acrfas012008* | *NACHT* | Immunity & Defense | NACHT, LRR and PYD domains |
| CMY2CNA | *Acrfas002029* | *PRF1* | Immunity & Defense | Perforin 1 |
| CMY2CNA | *Acrfas010214* | *RNF213* | Immunity & Defense | E3 ubiquitin-protein ligase RNF213 |
| CMY2CNA | *Acrfas022406* | *SEMA7A* | Immunity & Defense | Semaphorin 7A |
| CMY2CNA | *Acrfas008333* | *STAB1* | Immunity & Defense | Stabilin 1 |
| CMY2CNA | *Acrfas023130* | *TRIM47* | Immunity & Defense | Tripartite motif containing 47 |
| CMY2CNA | *Acrfas023121* | *TRIM47* | Immunity & Defense | Tripartite motif containing 47 |
| CMY2CNA | *Acrfas018890* | *CACNA2D1* | Osmoregulation & Ion Transport | Ca2+ channel alpha-2/delta subunit |
| CMY2CNA | *Acrfas008082* | *CNG-K+ Channel* | Osmoregulation & Ion Transport | Cyclic nucleotide-gated K+ channel |
| CMY2CNA | *Acrfas027446* | *MICU3* | Osmoregulation & Ion Transport | Mitochondrial calcium uptake |
| CMY2CNA | *Acrfas018035* | *SLC35F3* | Osmoregulation & Ion Transport | Solute carrier family 35 |
| CMY2CNA | *Acrfas026067* | *SLC6A19* | Osmoregulation & Ion Transport | Neutral amino acid transporter |
| CMY2CNA | *Acrfas027861* | *STC2* | Osmoregulation & Ion Transport | Stanniocalcin 2 |
| CMY2CNA | *Acrfas004393* | *ACTB* | Muscle, Cytoskeleton & Adhesion | Actin Beta |
| CMY2CNA | *Acrfas007949* | *Adhesion-like* | Muscle, Cytoskeleton & Adhesion | Homophilic cell adhesion |
| CMY2CNA | *Acrfas024764* | *DNAH12* | Muscle, Cytoskeleton & Adhesion | Dynein heavy chain 12 |
| CMY2CNA | *Acrfas015256* | *KIF6* | Muscle, Cytoskeleton & Adhesion | Kinesin family member 6 |
| CMY2CNA | *Acrfas013434* | *MYH-Fast* | Muscle, Cytoskeleton & Adhesion | Myosin heavy chain, fast |
| CMY2CNA | *Acrfas013436* | *MYH-Fast* | Muscle, Cytoskeleton & Adhesion | Myosin heavy chain, fast |
| CMY2CNA | *Acrfas028189* | *MYOT* | Muscle, Cytoskeleton & Adhesion | Myotilin (Z-line) |
| CMY2CNA | *Acrfas022887* | *NCAM2* | Muscle, Cytoskeleton & Adhesion | Neural cell adhesion molecule 2 |
| CMY2CNA | *Acrfas022505* | *NCAM2* | Muscle, Cytoskeleton & Adhesion | Neural cell adhesion molecule 2 |
| CMY2CNA | *Acrfas027643* | *PCDH* | Muscle, Cytoskeleton & Adhesion | Protocadherin |
| CMY2CNA | *Acrfas027642* | *PCDH* | Muscle, Cytoskeleton & Adhesion | Protocadherin |
| CMY2CNA | *Acrfas027644* | *PCDH* | Muscle, Cytoskeleton & Adhesion | Protocadherin |
| CMY2CNA | *Acrfas027641* | *PCDH* | Muscle, Cytoskeleton & Adhesion | Protocadherin |
| CMY2CNA | *Acrfas015209* | *REEP1* | Muscle, Cytoskeleton & Adhesion | Receptor accessory protein 1 |
| CMY2CNA | *Acrfas021418* | *SYNE2* | Muscle, Cytoskeleton & Adhesion | Spectrin repeat containing |
| CMY2CNA | *Acrfas009847* | *TJP2* | Muscle, Cytoskeleton & Adhesion | Tight junction protein 2 |
| CMY2CNA | *Acrfas026686* | *ABO* | Metabolism & Mitochondria | Glycosyltransferase (ABO) |
| CMY2CNA | *Acrfas011762* | *CAPN2* | Metabolism & Mitochondria | Calpain 2 |
| CMY2CNA | *Acrfas027442* | *MTMR10* | Metabolism & Mitochondria | Myotubularin related protein 10 |
| CMY2CNA | *Acrfas008795* | *OS9* | Metabolism & Mitochondria | OS9 (ER lectin) |
| CMY2CNA | *Acrfas013095* | *PSMD12* | Metabolism & Mitochondria | Proteasome subunit |
| CMY2CNA | *Acrfas019651* | *XRCC6BP1* | Metabolism & Mitochondria | ATP23 (Mitochondrial protease) |
| CMY2CNA | *Acrfas027870* | *TENM2* | Nervous & Sensory System | Teneurin transmembrane protein 2 |
| CMY2CNA | *Acrfas010805* | *TMEM131* | Nervous & Sensory System | Transmembrane protein 131 |
| CMY2CNA | *Acrfas018907* | *UNC80* | Nervous & Sensory System | UNC80 homolog |
| CMY2CNA | *Acrfas027562* | *ADAMTS2* | Cellular Processes & Regulation | ADAM metallopeptidase |
| CMY2CNA | *Acrfas014167* | *AP3B1* | Cellular Processes & Regulation | Adaptor-related protein complex 3 |
| CMY2CNA | *Acrfas006884* | *AP5M1* | Cellular Processes & Regulation | Adaptor related protein complex 5 |
| CMY2CNA | *Acrfas003986* | *BLM* | Cellular Processes & Regulation | Bloom syndrome RecQ helicase |
| CMY2CNA | *Acrfas008606* | *CDK18* | Cellular Processes & Regulation | Cyclin-dependent kinase 18 |
| CMY2CNA | *Acrfas016551* | *ETV1* | Cellular Processes & Regulation | Ets variant gene 1 |
| CMY2CNA | *Acrfas027432* | *FAM189A1* | Cellular Processes & Regulation | Family with sequence similarity 189 |
| CMY2CNA | *Acrfas008362* | *FBXO42* | Cellular Processes & Regulation | F-box protein 42 |
| CMY2CNA | *Acrfas010291* | *FBXW8* | Cellular Processes & Regulation | F-box and WD repeat domain |
| CMY2CNA | *Acrfas016650* | *GSG2* | Cellular Processes & Regulation | Germ cell associated 2 (Haspin) |
| CMY2CNA | *Acrfas021452* | *GTF3C2* | Cellular Processes & Regulation | General transcription factor 3C |
| CMY2CNA | *Acrfas011577* | *LONRF2* | Cellular Processes & Regulation | LON peptidase N-terminal |
| CMY2CNA | *Acrfas003746* | *LRRK2* | Cellular Processes & Regulation | Leucine-rich repeat kinase 2 |
| CMY2CNA | *Acrfas005626* | *MAP4K1* | Cellular Processes & Regulation | Mitogen-activated protein kinase |
| CMY2CNA | *Acrfas016147* | *MATK* | Cellular Processes & Regulation | Megakaryocyte-associated tyrosine kinase |
| CMY2CNA | *Acrfas003154* | *MON2* | Cellular Processes & Regulation | MON2 homolog |
| CMY2CNA | *Acrfas024738* | *NUGGC* | Cellular Processes & Regulation | Nuclear GTPase |
| CMY2CNA | *Acrfas024737* | *NUGGC* | Cellular Processes & Regulation | Nuclear GTPase |
| CMY2CNA | *Acrfas003311* | *NUP205* | Cellular Processes & Regulation | Nucleoporin 205 |
| CMY2CNA | *Acrfas013315* | *PPEF2* | Cellular Processes & Regulation | Protein phosphatase EF-hand |
| CMY2CNA | *Acrfas004287* | *PPP1R9B* | Cellular Processes & Regulation | Protein phosphatase 1 regulatory |
| CMY2CNA | *Acrfas002400* | *PTCHD3* | Cellular Processes & Regulation | Patched domain containing 3 |
| CMY2CNA | *Acrfas010807* | *PTGFRN* | Cellular Processes & Regulation | Prostaglandin F2 receptor |
| CMY2CNA | *Acrfas009654* | *RAD9B* | Cellular Processes & Regulation | Cell cycle checkpoint control |
| CMY2CNA | *Acrfas023951* | *RANBP2* | Cellular Processes & Regulation | RAN binding protein 2 |
| CMY2CNA | *Acrfas007035* | *RB1* | Cellular Processes & Regulation | Retinoblastoma 1 |
| CMY2CNA | *Acrfas018744* | *RGS17* | Cellular Processes & Regulation | Regulator of G-protein signaling |
| CMY2CNA | *Acrfas019070* | *RPS16* | Cellular Processes & Regulation | Ribosomal protein S16 |
| CMY2CNA | *Acrfas023092* | *SMC1B* | Cellular Processes & Regulation | Structural maintenance of chromosomes |
| CMY2CNA | *Acrfas026087* | *SNX13* | Cellular Processes & Regulation | Sorting nexin 13 |
| CMY2CNA | *Acrfas027985* | *SNX25* | Cellular Processes & Regulation | Sorting nexin 25 |
| CMY2CNA | *Acrfas010527* | *STK* | Cellular Processes & Regulation | Serine threonine-protein kinase |
| CMY2CNA | *Acrfas010862* | *SUMO1* | Cellular Processes & Regulation | SUMO1 |
| CMY2CNA | *Acrfas004123* | *SYNGR1* | Cellular Processes & Regulation | Synaptogyrin 1 |
| CMY2CNA | *Acrfas003919* | *TLE3* | Cellular Processes & Regulation | Transducin-like enhancer of split 3 |
| CMY2CNA | *Acrfas010730* | *TRHDE* | Cellular Processes & Regulation | Thyrotropin-releasing hormone |
| CMY2CNA | *Acrfas006968* | *USP28* | Cellular Processes & Regulation | Ubiquitin specific peptidase 28 |
| CMY2CNA | *Acrfas027436* | *C15orf43* | Others/Unknown | Bouquet formation protein 2 |
| CMY2CNA | *Acrfas010768* | *CCDC81* | Others/Unknown | Coiled-coil domain containing 81 |
| CMY2CNA | *Acrfas013142* | *DNA-bind* | Others/Unknown | DNA binding |
| CMY2CNA | *Acrfas025191* | *DUF4371* | Others/Unknown | Domain of unknown function |
| CMY2CNA | *Acrfas027828* | *DUF4371* | Others/Unknown | Domain of unknown function |
| CMY2CNA | *Acrfas006989* | *Endonuclease* | Others/Unknown | DDE superfamily endonuclease |
| CMY2CNA | *Acrfas022234* | *FSD2* | Others/Unknown | Fibronectin type III and SPRY |
| CMY2CNA | *Acrfas014928* | *HEATR4* | Others/Unknown | HEAT repeat containing 4 |
| CMY2CNA | *Acrfas013137* | *MAMDC4* | Others/Unknown | MAM domain containing 4 |
| CMY2CNA | *Acrfas023091* | *RT* | Others/Unknown | Reverse transcriptase |
| CMY2CNA | *Acrfas020328* | *Si-ch73* | Others/Unknown | Si ch73-132f6.5 |
| CMY2CNA | *Acrfas027975* | *Sugar-trans* | Others/Unknown | Sugar transporter (putative) |
| CMY2CNA | *Acrfas026267* | *Transposition* | Others/Unknown | RNA-mediated transposition |
| CMY2CNA | *Acrfas013186* | *Transposon* | Others/Unknown | RNA-mediated transposition |
| CMY2CNA | *Acrfas013574* | *Uncharacterized* | Others/Unknown | Transposition, RNA-mediated |
| CMY2CNA | *Acrfas018066* | *Uncharacterized* | Others/Unknown | - |
| CMY2CNA | *Acrfas009632* | *Uncharacterized* | Others/Unknown | - |
| CMY2CNA | *Acrfas021450* | *Uncharacterized* | Others/Unknown | - |
| CMY2CNA | *Acrfas019071* | *Zgc* | Others/Unknown | Zgc 113227 |
| CGR2CNA | *Acrfas019053* | *TLR* | Immunity & Defense | Toll-like receptor |
| CGR2CNA | *Acrfas025943* | *CLCN1* | Osmoregulation & Ion Transport | Chloride channel 1 |
| CGR2CNA | *Acrfas016585* | *FFAR* | Osmoregulation & Ion Transport | Free fatty acid receptor |
| CGR2CNA | *Acrfas024114* | *KCNAB2* | Osmoregulation & Ion Transport | Potassium channel subunit |
| CGR2CNA | *Acrfas027861* | *STC2* | Osmoregulation & Ion Transport | Stanniocalcin 2 |
| CGR2CNA | *Acrfas008111* | *MTUS2* | Muscle, Cytoskeleton & Adhesion | Microtubule associated tumor suppressor |
| CGR2CNA | *Acrfas021418* | *SYNE2* | Muscle, Cytoskeleton & Adhesion | Spectrin repeat containing |
| CGR2CNA | *Acrfas019299* | *FGFR1OP2* | Nervous & Sensory System | FGFR1 oncogene partner 2 |
| CGR2CNA | *Acrfas003454* | *NRCAM* | Nervous & Sensory System | Neuronal cell adhesion molecule |
| CGR2CNA | *Acrfas003538* | *SOX6* | Nervous & Sensory System | Transcription factor SOX-6 |
| CGR2CNA | *Acrfas001964* | *WDR48* | Cellular Processes & Regulation | WD repeat domain 48 |
| CGR2CNA | *Acrfas007737* | *TMEM179B* | Others/Unknown | Transmembrane protein 179B |
| CNA2CGR | *Acrfas022987* | *GTPase IMAP* | Immunity & Defense | GTPase IMAP family member |
| CNA2CGR | *Acrfas011215* | *IQCA1* | Muscle, Cytoskeleton & Adhesion | IQ motif containing with AAA domain |
| CNA2CGR | *Acrfas017946* | *AIFM2* | Metabolism & Mitochondria | Apoptosis-inducing factor 2 |
| CNA2CMY | *Acrfas007909* | *ATG* | Metabolism & Mitochondria | Autophagy-related protein |
| CNA2CMY | *Acrfas013867* | *PSMB7* | Metabolism & Mitochondria | Proteasome subunit beta 7 |
| CNA2CMY | *Acrfas008565* | *Ssu-2* | Metabolism & Mitochondria | Ssu-2 homolog |
| CNA2CMY | *Acrfas022472* | *PPIP5K1* | Cellular Processes & Regulation | Histidine acid phosphatase |
| CNA2CMY | *Acrfas003012* | *AARS* | Metabolism & Mitochondria | Alanyl-tRNA synthetase (Protein synthesis) |
| CNA2CMY | *Acrfas016171* | *TMC6* | Nervous & Sensory System | Transmembrane channel-like |
| CNA2CMY | *Acrfas019793* | *TRIM7* | Cellular Processes & Regulation | E3 ubiquitin-protein ligase |
| CNA2CMY | *Acrfas021564* | *MTHFD1* | Metabolism & Mitochondria | Methylenetetrahydrofolate dehydrogenase (NADP dependent) |
| CNA2CMY | *Acrfas020510* | *DNA-bind* | Cellular Processes & Regulation | DNA binding |
